# From Light to Acetate: How Trophic Conditions Shape Growth and Cell Cycle Progression in *Chlamydomonas reinhardtii*

**DOI:** 10.64898/2026.03.29.715089

**Authors:** Rabinder Singh, Firas Louis, Sien Audoor, P. V. Sijil, Martín Mora-García, Bipasha Bhattacharjee, Kateřina Bišová

## Abstract

The unicellular green alga *Chlamydomonas reinhardtii* provides a tractable model for investigating how carbon availability influences metabolic organization and cell-cycle control in photosynthetic eukaryotes. Its capacity for autotrophic (light, CO), mixotrophic (light, CO, acetate), and heterotrophic (acetate, dark) growth enables systematic analysis of trophic-state-dependent regulation. We performed comparative transcriptomic analyses of strain 21gr grown under these three regimes at 30 °C. Mixotrophy resulted in the highest biomass accumulation and was associated with earlier cell-cycle commitment compared with autotrophy, whereas heterotrophy displayed delayed commitment and reduced growth. Transcriptomic profiling revealed coordinated upregulation of central carbon metabolic pathways under mixotrophy, including photorespiration, glycolysis, the oxidative pentose phosphate pathway, and tricarboxylic acid cycle functions, consistent with enhanced carbon flux and biosynthetic capacity. In contrast, heterotrophy preferentially induced acetate assimilation and glyoxylate cycle genes and was accompanied by elevated expression of cell-cycle regulators, including the CDK-inhibitory kinase WEE1. Together, these findings indicate that trophic mode modulates the coupling between carbon metabolism and cell-cycle progression, with mixotrophy supporting integrated metabolic and proliferative activity, whereas heterotrophy is associated with delayed cell-cycle timing and transcriptional signatures of metabolic adjustment.

## Introduction

The unicellular green alga *Chlamydomonas reinhardtii* is a genetically tractable model organism for investigating eukaryotic photosynthesis (Dupuis and Merchant, 2023), cellular metabolism (Strenkert et al., 2019), and the cell cycle (Bišová and Zachleder, 2014). *C. reinhardtii* shares key pathways with other algae and land plants (Fauser et al., 2022) and offers the advantages of a unicellular model with a fully sequenced haploid genome and robust genetic tools (Merchant et al., 2007). It can grow photoautotrophically using light as the sole energy source for carbon dioxide fixation, heterotrophically by utilizing acetate as a carbon and energy source, or mixotrophically by combining both light and acetate (Boyle and Morgan, 2009; Füßl et al., 2022; Hoober, 1989). It grows easily at large scale and synchronizes with light/dark cycles, making it an ideal system for studying gene expression and cell division patterns (Strenkert et al., 2019). Despite significant progress, there remains a substantial gap in our understanding of the biochemistry and regulatory mechanisms governing metabolic processes in *C. reinhardtii*, especially regarding how the cells reprogram their physiology in response to varying energy and carbon sources. Each trophic mode imposes distinct metabolic demands, influencing key cellular processes such as glycolysis/gluconeogenesis, the tricarboxylic acid (TCA) cycle, amino acid biosynthesis, energy storage, and cell cycle progression (Füßl et al., 2022).

To assimilate C compounds such as acetate, C. reinhardtii uses an ATP-dependent entry into the glyoxylate cycle, generating C intermediates that can subsequently be converted into amino acids or soluble carbohydrates (Lauersen et al., 2016). Acetate suppresses photosynthesis while enhancing respiration under light and air bubbling, potentially through increased alternative oxidase activity (Endo and Asada, 1996; Fett and Coleman, 1994; Weger et al., 1990). Acetate availability is linked to altered expression of nuclear-encoded, chloroplast-targeted light-harvesting proteins (LHCII chlorophyll a/b–binding proteins), with acetate promoting their accumulation in the dark and attenuating expression during illumination (Kindle, 1987). The synergistic effect of acetate, light, and inorganic carbon attenuates the increased expression of isocitrate lyase, the essential glyoxylate cycle enzyme required for acetate assimilation (Kindle, 1987; Martínez-Rivas and Vega, 1993). Consistent with light-dependent metabolic reprogramming, *C. reinhardtii* tends to route acetate through glyoxylate cycle–linked metabolism under light, whereas in darkness mitochondrial respiration increases and TCA cycle activity becomes comparatively more active (Heifetz et al., 2000). An assimilatory kinetics study on heterotrophy and mixotrophy in *C. reinhardtii* showed complete consumption of 20 mM acetate within the first 12 hours in heterotrophy. The kinetics were slower in mixotrophy, suggesting competing entry of carbon via CO fixation during mixotrophic growth (Singh et al., 2014). A transcriptomic study under photoautotrophic and heterotrophic conditions showed distinct gene expression patterns, with genes related to glycolysis, the TCA cycle, and oxidative phosphorylation upregulated during heterotrophic growth, suggesting an efficient supply of energy and carbon skeletons for rapid growth of C. reinhardtii (Chen et al., 2024). This aligns with metabolic flux balance analyses showing that C. reinhardtii utilizes distinct metabolic pathways depending on its growth conditions (Boyle and Morgan, 2009). A recent isotope-assisted metabolic flux analysis indicated that acetate induces a coordinated rewiring of central carbon metabolism, promoting carbon conservation via activation of the glyoxylate cycle and repression of gluconeogenesis. Despite partial inhibition of photosynthesis, acetate supplementation supported higher growth rates in mixotrophy compared to autotrophic growth (Koley et al., 2026). Rapid metabolic adaptation to acetate may also be facilitated by post-translational protein modifications, which act as a reversible switch enabling proteins to change their activities, functions, or localizations in the cell (Castaño-Cerezo et al., 2014; Kaur et al., 2021).

*C. reinhardtii* divides by multiple fission, a mechanism in which the cell increases its size manyfold (routinely more than eightfold) during a prolonged G1 phase. This growth phase is followed by multiple rounds (*n*) of alternating S/M phases, yielding numerous daughter cells (2*^n^*, usually 4–16). This strategy allows the cells to separate growth and division both temporally and functionally (Bišová and Zachleder, 2014; Cross and Umen, 2015). During the G1 phase, the cell reaches a control point, the commitment point (CP), where it becomes irreversibly committed to completing a division sequence even if conditions change (e.g., transfer from light to dark). The CP is conceptually similar to START in yeast and the restriction point in animal cells (Bišová and Zachleder, 2014). To reach the CP, the cell must grow until it attains a critical size. Thus, CP attainment is growth-dependent, and faster growth (at higher light and more favorable temperatures) leads to earlier CP. The critical cell size remains constant over a wide range of light conditions (Vítová et al., 2011). Prolonged growth beyond the CP leads to attainment of additional CPs; commonly, three CPs are reached within one cell cycle, resulting in the production of eight daughter cells. A mitotic sizer couples mother cell size to division number (*n*) so that daughter size distributions are uniform regardless of mother size distributions (Liu et al., 2023). The combination of two (interlinked) sizers, a CP sizer and a mitotic sizer, ensures uniform CP and daughter cell sizes and makes these sizes an important characteristic of cell behavior.

Here, we studied cell cycle progression in synchronized cultures under different trophic conditions, reasoning that trophic conditions are a fundamental aspect of natural ecosystems and cells should be able to respond accordingly to acclimate to environmental changes. This study provides a direct comparison of the effects of the three trophic modes, under the same growth conditions, on cell growth and cell cycle progression, offering insights into optimal growth conditions and cell acclimation to changing environments. Our main goal was to analyze CP attainment under these different trophic conditions, hypothesizing that CP attainment is growth-dependent but also serves as the main ON switch for cell cycle entry, thereby translating different metabolic inputs into entry or non-entry outcomes. To this end, we sampled at biologically defined time points before and after CP attainment to identify components and mechanisms shared by or distinct to CP attainment under different trophic conditions. This study focuses on the distinct features; the common ones will be described elsewhere. The timing of the sampling points also allowed us to analyze early phases after the trophic switch. We present an integrated analysis of gene expression, cell cycle organization, photosynthetic performance, and energy reserves in synchronized cultures grown under autotrophic (AUTO), mixotrophic (MIXO), and heterotrophic (HETERO) conditions. Of the three cultures, MIXO exhibited high metabolic activity, cellular respiration, carbohydrate and lipid metabolism, and a highly active TCA cycle. HETERO conditions showed delayed DNA replication, increased DNA damage, and elevated expression of the CDK inhibitory kinase WEE1, which was not observed in the other two conditions. The data provide comprehensive insights into the metabolic and regulatory adaptations of C. reinhardtii under varying trophic conditions and demonstrate the complex interplay between carbon availability, energy metabolism, and cell cycle progression.

## Materials and methods

### Growth conditions

*Chlamydomonas reinhardtii 21 gr* (CC–1690, wild type; Chlamydomonas Resource Center, St. Paul, Minnesota, USA, http://www.chlamy.org, accessed on 27 July 2025) was grown in tubular glass vessels (final volume 300 ml) in a high salt mineral medium (HSM) (Harris, 2001), at 30 ◦C, with air-CO_2_ (2% v/v) aeration. The vessels were illuminated from one side by fluorescent tubes (OSRAM DULUX L55 W/950 Daylight, Milano, Italy), the light intensity at the culture vessel surface was 500 μmol·m^−2^·s^−1^ of photosynthetically active radiation (PAR).

To synchronize population growth, alternating cycles of 12 hour (h) light and 12 h dark were applied to the cultures being diluted to a constant cell number at the beginning of each light period (Coleman, 1982; Hlavová et al., 2016). At the beginning of the experiment, the synchronized cell population was diluted to a starting concentration of ∼1×10^6^ cell. mL^-1^ in either HSM (AUTO) or TAP medium (MIXO, HETERO) (Burgess et al., 2016; Gorman and Levine, 1965) and grown at 30 °C under continuous light of 500 μmol·m^−2^·s^−1^ of PAR for AUTO and MIXO, and in dark for HETERO. All the cultures were bubbled by 2% (v/v) (2,000 ppm) CO_2_ in air. The experiments were run as biological triplicates, and the cultures were sampled at regular intervals (either hourly or bi-hourly) (Fig. 1).

**Figure 1.**
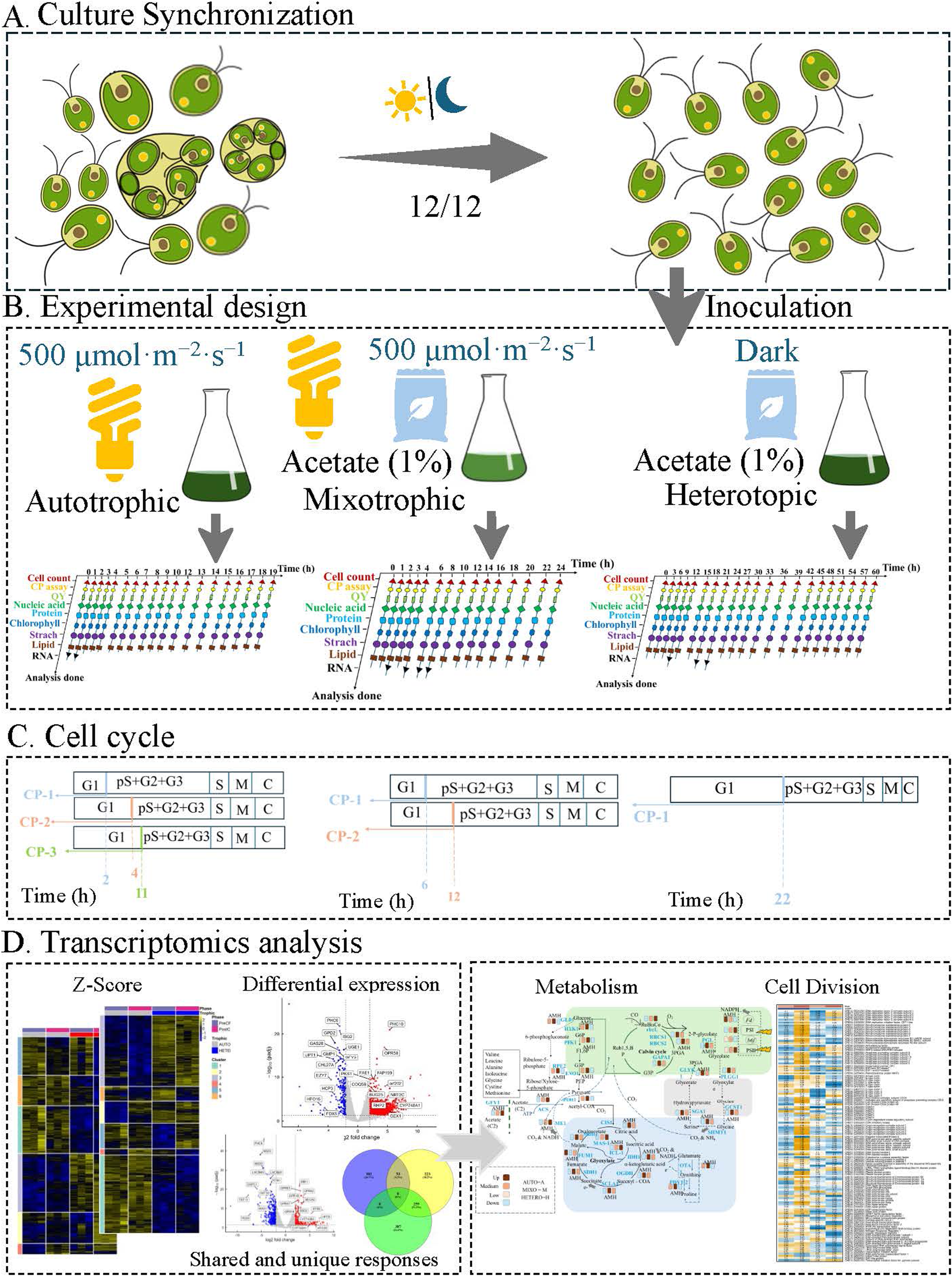
Schematic representation of the experimental design. A) *Chlamydomonas reinhardtii* cultures were synchronized using 12 h light/12 h dark cycles at 30 °C and an irradiance of 500 µmol m ² s ¹. B) For the experiment, synchronized cells were diluted to approximately 1 × 10 cells mL ¹ in HSM (AUTO) or TAP (MIXO, HETERO) and grown at 30 °C under continuous light (500 µmol m ² s ¹ PAR), bubbled with 2% CO in air. Experiments were performed in biological triplicates with hourly or bi-hourly sampling for different analyses. C) Representation of cell cycle events in each trophic condition. D) Schematic representation of the transcriptomics analysis workflow.

### Cell cycle analyses and cellular division evaluation

The attainment of CPs was assessed (bi-)hourly in aliquots (1 mL) of the culture placed on HSM plates and incubated in the dark at 30 °C until cell division was completed in the parental culture (Kselíková et al., 2022). The plates were then examined directly under a light microscope (magnification 10 × 40). For each sample, the number of large single undivided mother cells and mother cells divided into 2, 4, or 8 daughter cells (at least 150 mother cells) were counted, and the cumulative percentage of each category was calculated and plotted against time. Mother and daughter cells can be easily distinguished by differences in cell size, and cell clumps can be distinguished by the specific morphology of the divided mother cells.

Cell division was assessed in samples that were taken (bi-)hourly, fixed with glutaraldehyde (final concentration 0.2%) and observed under a light microscope (magnification 10 × 40). The cumulative percentage of undivided mother cells or mother cells divided into 2, 4, or 8 daughter cells were calculated and plotted against time.

Cell culture doubling time (*Td*) was calculated using the formula:

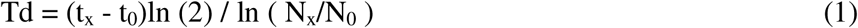

At the start of the experiment (*t_0_*), *N_0_* represents the initial number of cells, while *N_x_* denotes the number of cells measured at the end of the experiment (time point *t_x_*) (Neumann et al., 2004). The time intervals are expressed in hours. Additionally, the cell division number was determined using the ratio *N_x_/N_0_*, where *N_0_* and *N_x_* correspond to the cell counts at the beginning and end of the experiment, respectively.

### Cell volume and dry mass

Cell volumes were measured with the Multisizer 4e (Beckman Coulter) in glutaraldehyde fixed samples (final concentration 0.2%) by diluting 50 μL of fixed cells in 10 mL 0.9% NaCl. The results were given as modal volume values of the cells in the sample. For determining daughter cell size, 10 mL aliquots from 9 h and 11 h of the cell cycle were aerated at 30 ◦C in the dark until division was complete (not more than 24 h from start of illumination), samples were fixed with glutaraldehyde and analyzed.

Biomass was separated from the medium by centrifugation of 5 mL of the cell suspension in pre-weighed microtubes at 3000× g for 5 min; the sediment was dried at 105 °C for at least 12 h (until stable weight) and weighed on an analytical balance (TE214S-0CE, Sartorius, Goettingen, Germany) (Brányiková et al., 2011).

### Quantum yield

The quantum yield of photosynthesis was measured as the F_V_/F_M_ ratio of chlorophyll fluorescence. Aliquots of 2 ml were taken from the culture and darkened for 30 min in 10 mm × 10 mm plastic cuvettes at room temperature. The cell suspension was mixed gently by inverting the cuvettes several times before measurement. Quantum yield was measured using an Aqua-Pen-C 100 (Photon Systems Instruments, Drasov, Czech Republic) set according to the manufacturer’s instructions (Genty et al., 1989).

### DNA, RNA and proteins quantification

Total nucleic acids were extracted according to (Wanka, 1962), as modified by (Lukavský et al., 1973). The DNA assay was carried out as described by (Decallonne and Weyns, 1976), with modifications of (Zachleder, 1984). The sediment remaining after nucleic acid extraction was quantified for protein content according to the procedure described by Lowry and Randall (Zachleder, 1984). The data are expressed in pg of DNA, RNA and protein content per cell normalized to the starting values.

### Statistical analysis

All experiments were performed in three biological replicates (n = 3). Presented results are the average and standard deviations from all three replicates. MS Excel 2010 (Microsoft) with Real Statistics Resource Pack (https://www.real-statistics.com/free-download/real-statistics-resource-pack/, accessed on 27 May 2025) was used to generate statistics. Comparison between experimental treatments was performed using the two-way ANOVA test and Tukey’s HSD test. A p-value < 0.05 was considered significant.

### RNA extraction and library preparation

RNA was isolated using the ZiXpress Viral DNA/RNA Extraction Kit (Zinexts; cat. no. ZP02201-192) on the ZiXpress robot (Zinexts) according to the manufacturer’s instructions. Contaminating DNA was removed by in-column DNase treatment (NEB; cat. no. M0303) according to the manufacturer’s instructions. RNA fragmentation and rRNA depletion were performed using the QIAseq FastSelect kit (QIAGEN; cat. no. 334315). Libraries were prepared using the NEBNext Ultra II Directional Library Prep Kit (NEB; cat. no. E7760) according to the manufacturer’s instructions. All purification steps were carried out using SPRIselect beads (Beckman Coulter; cat. no. B23319). Library quality was assessed by agarose gel electrophoresis. The libraries were subsequently sequenced by Novogene (paired end, 150 bp; PE150) on an Illumina platform.

### Transcriptome data analysis

Raw sequencing reads were assessed for quality using FastQC (Mielczarek et al., 2023). Adapters were removed from and the low-quality reads and cleaned using fastp (Chen, 2023). Cleaned reads were mapped to indexed *Chlamydomonas reinhardtii* CC-4532 v6.1 genome and transcriptome (https://phytozome-next.jgi.doe.gov/) (Craig et al., 2022), using Salmon (version 1.10.2) (Patro et al., 2017). The Salmon quantification files were made readable using tximport (Mielczarek et al., 2023). PCA of the transcriptome was performed separately on each comparison using the R package PCA tools (v2.16.0) on mRNAs with a minimum average FPKM (17, 369 genes). The count data were tested for differential expressions using edgeR (v4.8.2) (Chen et al., 2025), with the GLM likelihood algorithm (Love et al., 2014). Significant differential expression was defined as |log2 fold-change | > 2, FDR < 0.05. Mean transcript abundances (as FPKMs) were Z-score normalized across different trophic conditions (AUTO VS MIXO, HETERO vs AUTO, HETERO vs MIXO). Heatmaps were generated using the R package pheatmap (version 1.0.13. https://CRAN.R-project.org/package=pheatmap). Functional enrichment analysis was performed using MapMan (Usadel et al., 2009) annotations for *Chlamydomonas reinhardtii* using *Chlamydomonas reinhardtii* CC-4532 v6.1. Common transcriptional expression patterns throughout AUTO, MIXO, and HETERO treatments, we performed by k-means clustering of their normalized expression patterns, implemented by pRocessomics R package (v.0.1.13, github.com/Valledor/pRocessomics)(Guerrero et al., 2024).

## Results

### Growth in heterotrophy is very slow

To directly compare the effect to the different trophic regimes on cell growth, metabolism, and cell cycle progression, we synchronized cultures of *C. reinhardtii* in minimal medium and exposed them to different trophic regimes for a duration of a single cell cycle (Fig. 1). The experiment started with cultures composed of daughter cells at concentration approximately 1×10^6^ cell·mL^-1^. The individual cultures were provided with CO_2_ alone (AUTO) or in combination with acetate (MIXO, HETERO) in light (500 µmol m^-2^ s^-1^, AUTO, MIXO) or in dark (HETERO). Cellular growth in algae is largely governed by the activity of organelles such as chloroplasts containing photosynthetic machinery. Chlorophyll *a* fluorescence is a quick and reliable indicator for detecting stress to photosynthesis responses in microalgae and serves as physiological indicator of cellular responses to varying environmental conditions (Minhas et al., 2016). To assess the efficiency of light energy conversion during photosynthesis, we measured the maximum quantum yield of photosystem II (Fv/Fm). All the values were within the normal physiological range, confirming no damaging effect of any trophic regime nor the light intensity used (Fig. 2A). AUTO and MIXO depicted similar patterns of PSII (Fv/Fm) with slight drop at 1 hour after the light onset, then improved to previous values. At the end of cultivation, the QY for the MIXO culture dropped slightly, but for all conditions it remained above ∼0.5 suggesting non-stress growth conditions and healthy photosynthetic machinery (Baker, 2008). Between the two light grown cultures there was no difference in chlorophyl accumulation, while the dark grown HETERO culture did not accumulate any further chlorophyl (Fig. 2B).

**Figure 2.**
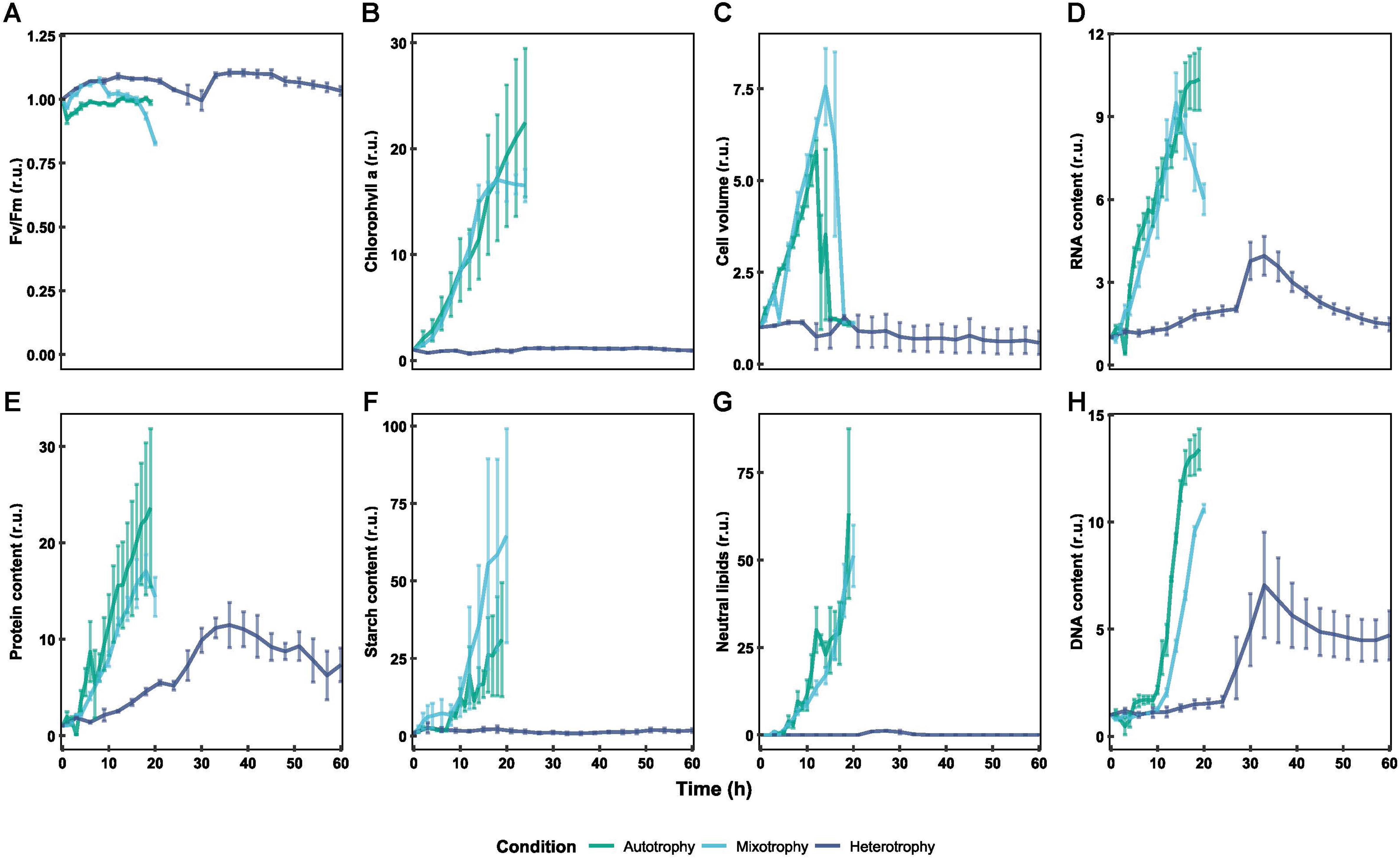
Growth of synchronized cultures of *Chlamydomonas reinhardtii* grown under different trophic regimes. A, maximum quantum efficiency of photosystem II (Fv/Fm); B, chlorophyll a; C, mean cell volume; D, RNA; E, protein; F, starch; G, neutral lipids; H, DNA. Except of Fv/Fm, all values were normalized to the first value in the dataset. All the experiment are represented as (n=3) biological replicates.

When assessing the growth, the MIXO cultures grew the fastest of the three (Fig. 2C). This was further confirmed by final dry cell weight (DCW) which was highest under MIXO, reaching 617.07±130.79 pg·cell ¹, followed by AUTO at 308.85±63.85 pg·cell ¹. HETERO cultures grew the slowest and accumulated the lowest biomass, 44.72±14.08 pg·cell ¹ (Fig. 2C). The cell volumes (µm^3^) ranged in AUTO between 56.73 µm^3^±1.00 (in young daughter cells at the beginning of the cell cycle) and 329.17 µm^3^±41.39 (mature mother cells at the 12^th^ h before division), i.e. increase about 5.8 fold (Fig. 2C). These large cells began dividing after 14 h, resulting in a decrease in cell volume to 56.73 µm^3^±2.07 and a concomitant increase in cell density. Similarly, in MIXO cell volumes ranged from 49.56 µm^3^ ± 5.49 in young daughter cells to 368.23 µm^3^±49.01 in mature mother cells at the 14^th^ h, i. e. increase about 7.4 fold. In HETERO, cell volumes didn’t change significantly, young daughter cells of size 52.04 µm^3^±1.99, remained same in size throughout experiment (Fig. 2C). Total RNA content increase (5.7-fold in MIXO, 10-fold in AUTO) did not match the extent of cell volume increases, suggesting different cell composition between the two growth modes (Fig. 2D). In HETERO cultures, the accumulation of total RNA was notably slow, and the RNA content increased only ∼1.49-fold (Fig. 2D). Similar pattern was also seen for the bulk protein content (Fig. 2E).

Starch accumulation was the highest in MIXO conditions, increasing more than 43-fold over a period of 20 hours. In AUTO, starch content increased about 33-fold within 19 hours. The lowest starch accumulation was observed in HETERO, with increase l.7-fold, i. e. less than two-fold effectively decreasing the cell starch content between the daughter cells at the beginning and end of the experiment (Fig. 2F). Lipid accumulation was similar in MIXO and AUTO, reaching about 50-fold increase within 20 hours. In HETERO, the lipid content was negligible at the threshold of detection, and no lipid accumulation was observed throughout the growth period (Fig. 2G).

### Cell Cycle Progression Differs between Growth Regimes

Synchronized culture in AUTO, sequentially attained three CPs, with midpoints at approximately 1.5, 5, and 11 h after the onset of light (Fig. 3A). The total cell cycle duration in AUTO was 19 hours. Similarly, MIXO attained three CPs, with midpoints at approximately 2, 4, and 10 h (Fig. 3B). While the first CP attainment was comparable between the two treatments, the additional CP attainments in MIXO were advanced compared to AUTO. In HETERO, CP attainment was delayed with only about 50 % of the cells passing the first CP at approximately 18 h. The second CP was passed only in about 30% of population (Fig. 3C). Consistent with the low commitment rate, only ∼50% of cells divided, producing two to four daughter cells. The critical sizes at which first CP was attained were comparable among all treatments (Table 1).

**Figure 3.**
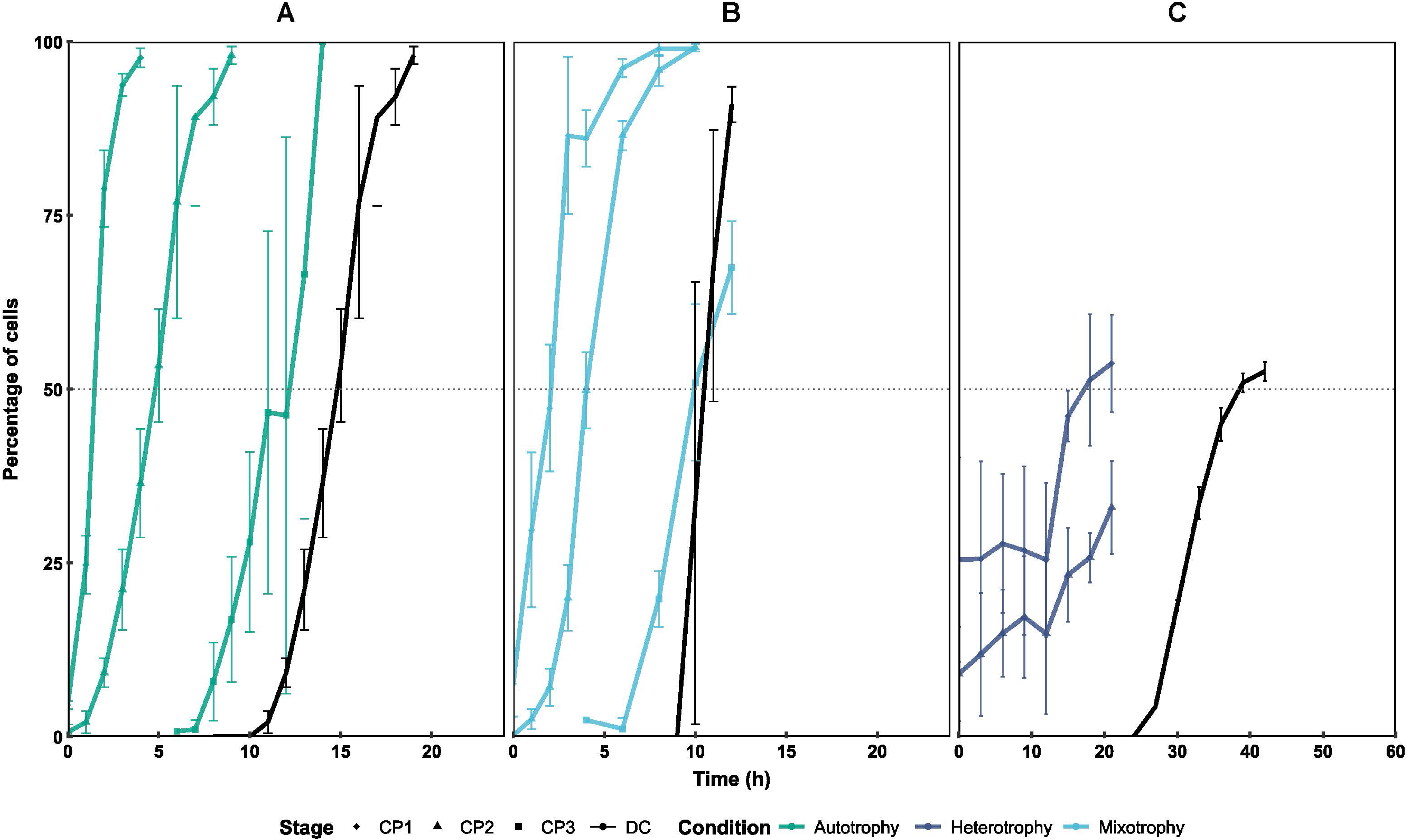
Cell cycle progression in a synchronized culture of *C. reinhardtii* grown under different trophic regimes. Time courses of individual commitment points (CP, colored lines) and daughter cell release (black lines) in autotrophy (A), mixotrophy (B), and heterotrophy (C). The colored curves show the cumulative percentage of cells that attained the CP for division into 2 (squares), 4 (circles), and 8 (triangles) daughter cells. The black lines show the cumulative percentage of cells that completed cell division. The gray dashed line depicts the midpoint of each event in the cell population. All experiments are represented as biological replicates (n = 3).

**Table 1.**
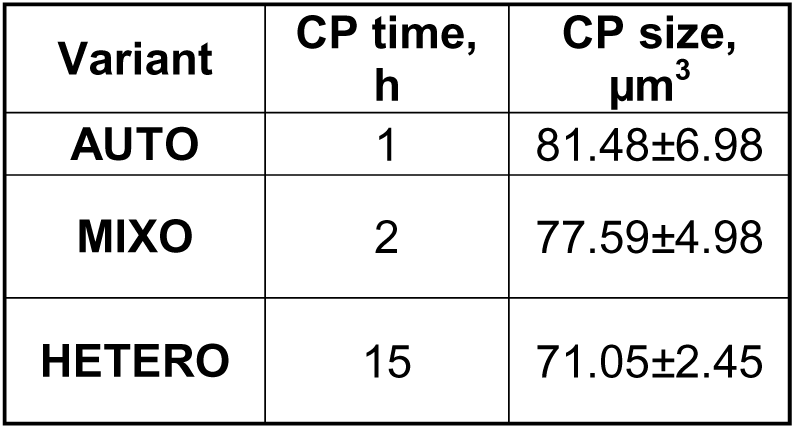
Time of reaching commitment point and critical cell size in *Chlamydomonas reinhardtii* under different trophic regimes.

S/M phase started by the 11^th^ to 12^th^ hour of the cell cycle in both AUTO and MIXO. It was completed by the 16^th^ hour in AUTO and 20^th^ hour in MIXO. During this time, DNA multiplied about 12-fold in AUTO and about 10-fold MIXO (Fig. 2H) corresponding to the production of about 8 daughters per mother cells. In HETERO, the total DNA rose about 6-fold over the 60-hour cultivation period (Fig. 2H).

### Distinct transcriptomic profile in heterotrophy

Due to differences in growth among the regimes, sampling them at the same time after the trophic switch would introduce additional variability, as samples would be affected by both the trophic regime and the growth rate. To avoid this, we sampled for transcriptomics at biologically equivalent timepoints: right before CP attainment (Pre-CP) and soon after CP attainment (Post-CP), that is, at times when cells in any condition reached critical size and were ready to enter the cell cycle (Fig. 1). The timing of Pre-CP and Post-CP sampling was determined independently for each trophic condition based on the commitment point assay, ensuring that samples corresponded to similar biological states despite differences in growth rates between AUTO, MIXO, and HETERO. While CP attainment in AUTO and MIXO conditions occurred at a similar timepoint early after cell cycle start, it was significantly delayed in HETERO cultures. As a result, the samples from AUTO and MIXO conditions were inherently closer both biologically and temporally. Given the rapid CP attainment in AUTO and MIXO, these samples reflected the immediate effects of the trophic switch. In contrast, samples from HETERO cultures reflected acclimation to the trophic switch. The sampling design thus captured (i) transcriptional responses associated with early metabolic reprogramming after the trophic switch (Pre-CP), and (ii) entry into cell division (Post-CP).

We detected approximately 17,369 genome-wide transcripts in all trophic regimes (AUTO, MIXO, and HETERO). Across the genome, RNA abundances varied depending on the trophic conditions. Principal component analysis (PCA) of the RNA-Seq data revealed that most transcriptomic variation was driven by differences in growth conditions, with only a smaller proportion attributable to the cell cycle stage (Fig. 4). Additionally, strong similarity was found between biological replicates, as demonstrated by Pearson correlation coefficients (*r*), confirming reproducibility (Supplementary Figure 1 and Supplementary dataset 1).

**Figure 4.**
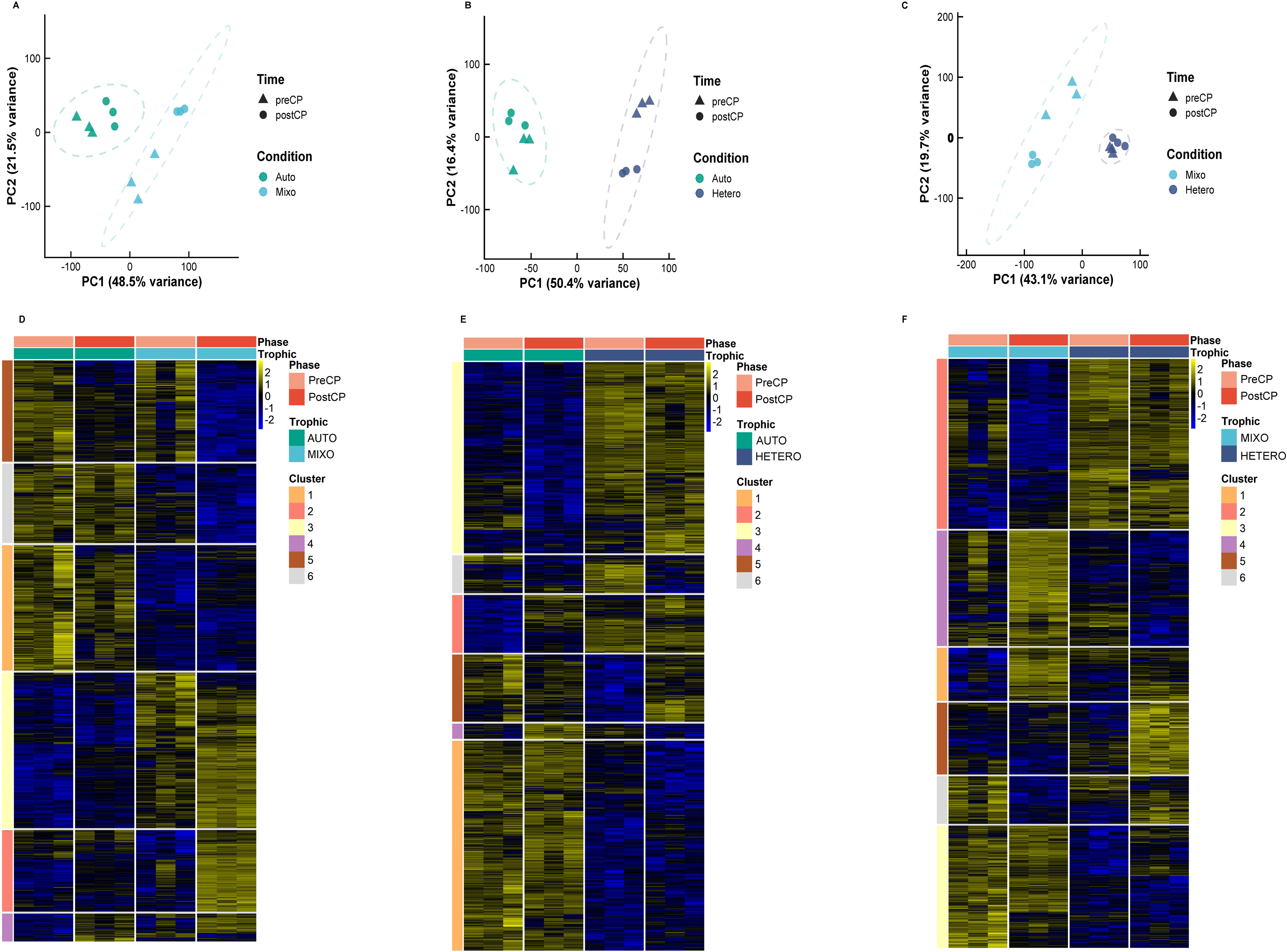
Principal Component Analysis (PCA) of genome-wide RNA abundances (FPKMs) across three biological replicates (n = 3). This analysis shows sample clustering based on gene expression profiles from RNA-seq data of C. reinhardtii. The first principal component (PC1) and the second principal component (PC2) are displayed. **A**, AUTO and MIXO, **B**, HETERO and AUTO, **C**, MIXO and HETERO. RNA expression was grouped into eight clusters by k-means clustering of their normalized expression patterns. Normalized values are the Z-scores of the mean mRNA abundance (FPKM) from three experimental replicates (n = 3) in **A**, AUTO and MIXO, **B**, HETERO and AUTO, and **C**, MIXO and HETERO.

We employed differential expression modeling to identify differentially expressed genes (DEGs) that were either overlapping or uniquely upregulated or downregulated in AUTO, MIXO, and HETERO treatments (Supplementary dataset 2, 3, and 4). Differentially expressed genes were considered significant if they had an adjusted p-value ≤ 0.05 and an absolute log2 fold change (|log2FC|) ≥ 2. The largest number of significant DEGs was identified between Post-CP MIXO vs. Post-CP HETERO, with approximately 6,098 significantly altered transcripts—3,328 upregulated and 2,770 downregulated (Fig. 5F; Supplementary dataset 4). This was followed by Post-CP AUTO vs. Post-CP HETERO, with about 5,017 significant transcripts, of which 2,455 were upregulated and 2,562 were downregulated (Fig. 5E; Supplementary Dataset 3). The difference between mixotrophic to heterotrophic growth and switch from autotrophic to heterotrophic growth was marked by strong oscillation in the expression of genes involved in heterotrophic metabolism, indicating that the absence of light and the presence of external carbon sources strongly induced heterotrophic metabolism. Furthermore, in Post-CP MIXO vs. Post-CP AUTO, approximately 4,575 significant DEGs were observed, with 2,079 upregulated and 1,196 downregulated (Fig. 5D; Supplementary Dataset 2). Additional shared and unique responses in these comparisons were also identified (Supplementary Dataset 2). To further analyze the biological processes underlying the global expression shifts, we performed pathway-level analyses. Below we outline several examples of metabolic pathways with distinct behavior under different trophic conditions. Differential expression was evaluated using log2 fold-change > 2 and FDR < 0.05, and expression patterns were also assessed by Z-score normalization.

**Figure 5.**
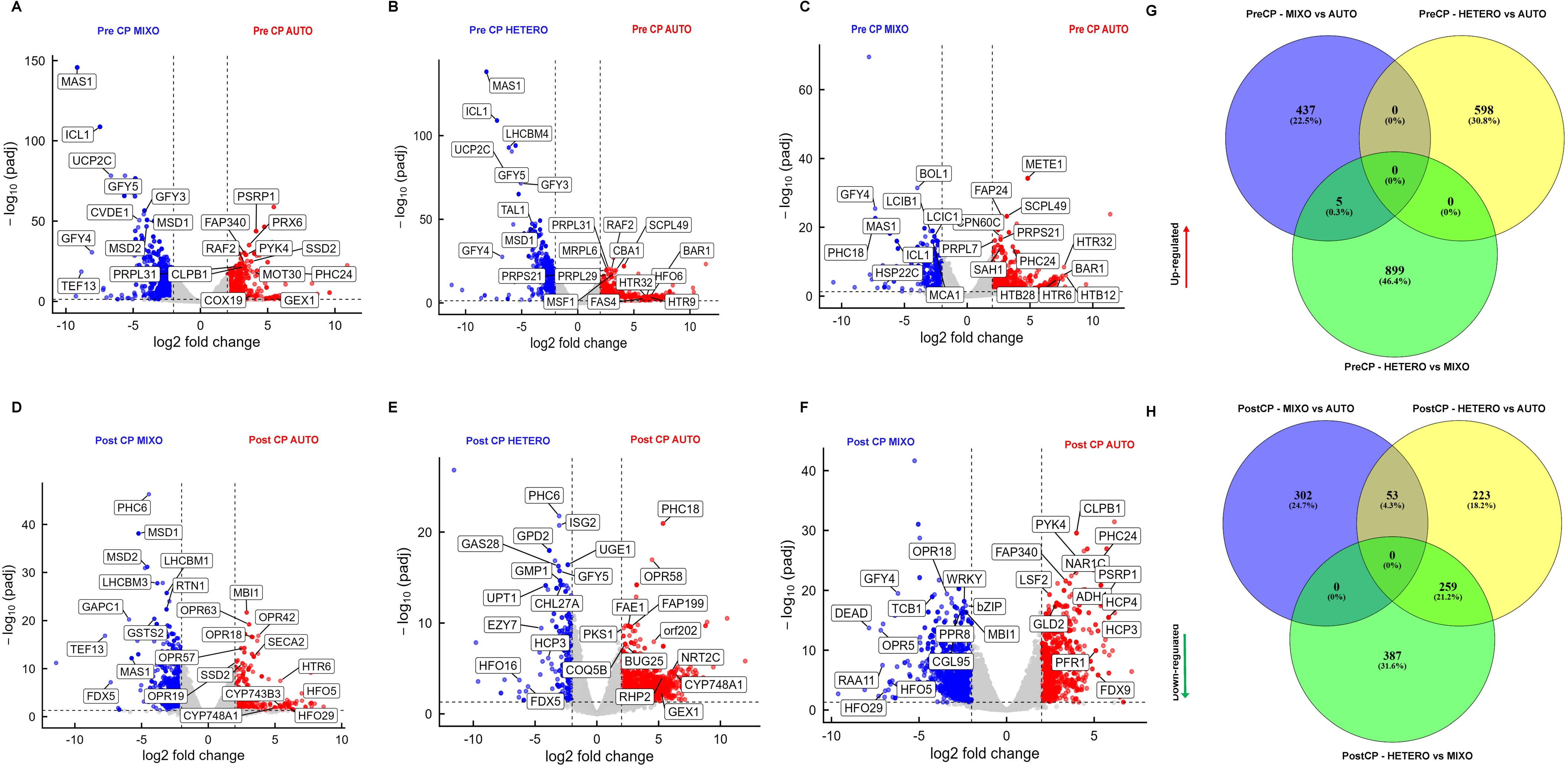
Differential gene expression analysis of C. reinhardtii under different growth conditions. Volcano plots show the correlation between log2(FC) values of genes and the corresponding –log (padj-values); only genes quantified in three biological replicates are included. **A**, Differential gene expression analysis: Pre-CP MIXO vs. Pre-CP AUTO; **B**, Pre-CP HETERO vs. Pre-CP AUTO; **C**, Pre-CP MIXO vs. Pre-CP HETERO; **D**, Post-CP MIXO vs. Post-CP AUTO; **E**, Post-CP HETERO vs. Post-CP AUTO; **F**, Post-CP MIXO vs. Post-CP HETERO. Venn diagrams of upregulated (**G**) and downregulated (**H**) genes show the shared and unique responses among the conditions.

#### Photosynthesis

In Post-CP MIXO compared with Post-CP AUTO, photosynthesis-related genes were upregulated, including the PSI light-harvesting protein LHCA4 (logFC = 2.19), LHCII chlorophyll a/b–binding proteins such as LHCBM1 (logFC = 3.19), and LHCII antenna subunits (logFC = 3.17) (Supplementary Dataset 2). In Post-CP HETERO compared with Post-CP AUTO, we found similar patterns, with LHCA4 (logFC = 3.43) and LHCA1 (logFC = 2.22874) upregulated in Post-CP HETERO (Supplementary Dataset 3). (Roach et al., 2013) demonstrated that acetate in TAP medium protects *C. reinhardtii* from photoinhibition by altering the PSII charge recombination reaction, thus lowering the production of singlet oxygen (^1^O) compared to autotrophy. In contrast, it has also been reported that under light conditions, acetate suppresses the light-induced increase in LHCII chlorophyll *a/b*–binding proteins from approximately 6 hours onward. Despite the difference in timing, our results may be consistent with this phenomenon as the growth conditions used in our experiments were supporting faster growth. Alternatively, we captured the initial induction phase rather than the later acetate-mediated repression (Kindle, 1987). In line with this, recent isotope-assisted metabolic flux analyses have shown that acetate decreases ^13^CO assimilation and Calvin-Benson-Bassham (CBB) cycle flux by approximately 31%, thereby partially suppressing photosynthesis. Despite this reduction, mixotrophic cells exhibit higher growth rates than autotrophic cells. These findings suggest that cells strategically downscale photosynthetic activity to reduce biosynthetic and energetic costs when acetate provides an efficient external carbon source (Koley et al., 2026).

Cytochrome b6/f complex genes (petD and petG) showed an increased trend in Post-CP MIXO based on z-score normalization and were downregulated in Post-CP HETERO (Fig. 4), indicating trophic state-specific remodeling of PSI light harvesting and the electron transport chain. Interestingly, we also found that, in Post-CP HETERO vs. Post-CP MIXO, the FdC-type ferredoxin electron carriers (FdC2 and Fd1/2/3) were upregulated in Post-CP MIXO; increased ferredoxin-dependent electron partitioning may support light-driven metabolism and biosynthetic demands during the post-CP-commitment phase in MIXO, as Fd completes the linear electron transport chain generating NADPH via ferredoxin–NADP reductase. In addition, upregulation of the thylakoid grana stacking regulatory factor (CRUT) was also found in Post-CP MIXO compared to Post-CP HETERO.

#### Acetate assimilation

In the Post-CP MIXO vs. Post-CP AUTO comparison, the acetate transporter *GFY1-5* (logFC = 2.91), acetyl-CoA synthase (*ACS1*) (logFC = 2.01), and phosphotransacetylase/phosphate acetyltransferase (*PAT*) (logFC = 2.02) were significantly upregulated (Fig. 5E), indicating enhanced acetate uptake and increased channeling of acetate-derived carbon into the acetyl-CoA pool and downstream central metabolism in mixotrophic cells. The involvement of GFY membrane proteins in acetate uptake has been demonstrated in *Escherichia coli* (Sá-Pessoa et al., 2013), *Citrobacter koseri* (Qiu et al., 2018), *Saccharomyces cerevisiae* (Paiva et al., 2004), and *Aspergillus nidulans* (Robellet et al., 2008), as well as their structural homology with *CrGFY1-5* (Durante et al., 2019). In addition, *GFY5* was strongly regulated in Post-CP HETERO relative to Post-CP AUTO (logFC = 4.86), consistent with the concomitant induction of *ACS2* (logFC = 3.04), a gateway enzyme that converts acetate to acetyl-CoA for entry into the glyoxylate cycle/TCA cycle and downstream central carbon metabolism (Johnson and Alric, 2013). Strikingly, when we compared Post-CP MIXO vs. Post-CP HETERO, *GFY4* (logFC = 6.20) was upregulated in Post-CP HETERO. This was also evident in the Z-score trend, with upregulation of *ACS* and *GFY1–5*, signifying enhanced acetate uptake and increased conversion of acetate into the acetyl-CoA pool to support carbon and energy metabolism during heterotrophic growth.

#### Photorespiration and carbon capture

Among the photorespiration-associated genes, serine glyoxylate aminotransferase (*SGA1*) was upregulated (logFC = 2.21) in Post-CP MIXO compared to Post-CP AUTO (Fig. 5D; Supplementary Dataset 2). Z-score trends showed coordinated upregulation of core photorespiratory components, including *AGT1*, *SHMT1*, *HPR1*, and the terminal enzyme *GLYK1*, consistent with increased photorespiratory recycling of 2-phosphoglycolate/glycolate to 3-phosphoglycerate to support redox homeostasis and carbon flux during mixotrophic growth (Fig. 6). A similar induction pattern was observed in Post-CP HETERO compared to Post-CP AUTO. Notably, although both conditions used acetate, the Post-CP MIXO vs Post-CP HETERO comparison showed higher Z-scores for *AGT1*, *SHMT1*, *HPR1*, and *GLYK1* in MIXO, indicating higher photorespiratory activity in MIXO than in HETERO (Fig. 5F).

**Figure 6.**
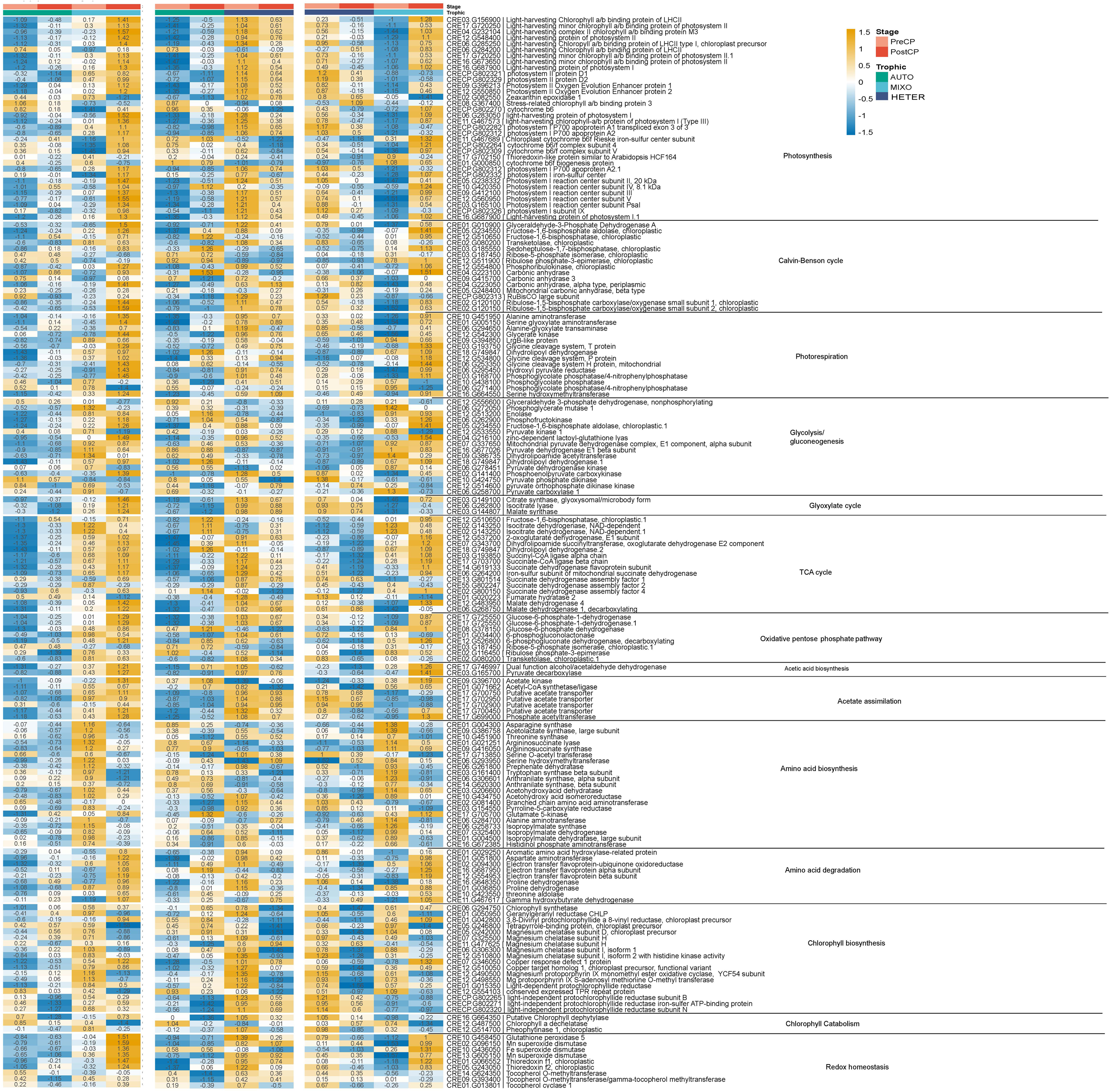
Z-score heatmaps show the expression of genes involved in central metabolic pathways in AUTO, MIXO, and HETERO under Pre-CP and Post-CP conditions. Colors indicate relative expression levels (yellow - increased; blue - decreased).

Genes involved in carbon capture and fixation were upregulated in Post-CP MIXO compared to Post-CP AUTO, including key CBB cycle components such as fructose-bisphosphate aldolase (*FBA2–3*), glyceraldehyde-3-phosphate dehydrogenase (*GAPA1*), sedoheptulose-1,7-bisphosphatase (*SBP1*), and transketolase (*TRK1*) (Fig. 6). In parallel, genes involved in central carbohydrate metabolism also showed higher expression, including the glycolytic genes enolase (*ENO1*), phosphofructokinase (*PFK1*), fructose-1,6-bisphosphate aldolase (*FBA3*), and pyruvate kinase (*PYK1*), consistent with elevated carbon flux to support post-CP commitment energy and biosynthetic demands under mixotrophic growth. In contrast, these genes were expressed at lower levels in Post-CP HETERO compared to both Post-CP MIXO and Post-CP AUTO, indicating that Post-CP MIXO maintains more photosynthetically driven, carbon assimilatory metabolic flux than Post-CP AUTO and Post-CP HETERO, despite the availability of acetate as an external carbon source. Furthermore, we observed repression of gluconeogenesis-associated genes in Post-CP MIXO compared to Post-CP AUTO; Post-CP HETERO showed a similar trend, including downregulation of pyruvate phosphate dikinase (*PPD1*). However, pyruvate carboxylase 1 (*PYC1*) was upregulated in Post-CP MIXO compared to both Post-CP HETERO and Post-CP AUTO. When acetyl-CoA is abundant, pyruvate carboxylase diverts pyruvate away from the TCA cycle, redirecting carbon toward different metabolic routes (Zhou et al., 2024).

#### Oxidative pentose phosphate pathway

Significant upregulation of Glucose-6-phosphate-1-dehydrogenase (*GLD1*) (logFC = 2.31) was found in Post-CP MIXO compared to Post-CP AUTO. Higher *GLD1* expression has been associated with increased oxidative pentose phosphate pathway (OPPP)–driven NADPH production, aligning with acetate-supported metabolism, biosynthetic demand, and/or redox balancing when carbon metabolism is elevated and/or photosynthetic reductant is constrained (Kruger and von Schaewen, 2003). Consistently, Z-score analysis revealed increased expression of multiple OPPP-associated genes in Post-CP MIXO, including *GLD1*, 6-phosphogluconolactonase (*PGL1*), ribose-5-phosphate isomerase (*RPI1*), transketolase (*TRK1*), and ribulose-phosphate 3-epimerase (*RPE2*). In contrast, *GLD1* was significantly upregulated (logFC = 2.29) in Post-CP AUTO compared to Post-CP HETERO (Supplementary Dataset 3), indicating reduced OPPP participation under heterotrophic post-CP commitment conditions relative to autotrophy and mixotrophy.

#### Glyoxylate and TCA pathway

Two key glyoxylate cycle genes, isocitrate lyase (*ICL1*) (logFC = 4.23) and malate synthase (*MAS1*) (logFC = 5.74), were significantly upregulated in Post-CP MIXO relative to Post-CP AUTO (Fig. 5D). Their induction was even more pronounced in Post-CP HETERO vs. Post-CP AUTO, reaching logFC values of 9.15 *(ICL1*) and 7.46 (*MAS1*), respectively (Fig. 5E). In addition, Z-score analysis showed higher expression of *ICL1* and *MAS1* in Post-CP HETERO compared to Post-CP MIXO, signifying higher glyoxylate activity in HETERO than in MIXO (Fig. 6).

Additionally, multiple TCA cycle genes – citrate synthase (*CIS2*), isocitrate dehydrogenase (*IDH2*), 2-oxoglutarate dehydrogenase (*OGD1*), succinyl-CoA ligase (*SCLA1*), fumarate hydratase (*FUM2*), and malate dehydrogenase (*MDH4*)—showed higher expression in Post-CP MIXO than in Post-CP HETERO. Notably, the succinate dehydrogenase assembly factor *SDHAF4* (logFC = 4.13) and *SCLA1* (logFC = 2.02) were strongly upregulated, supporting a more active TCA-linked respiratory metabolism under mixotrophic growth (Fig. 6).

#### Amino acid metabolism and redox homeostasis

Ornithine transaminase (*OTA1*; logFC = 2.40) and glycerol-3-phosphate dehydrogenase (*GPD2*; logFC = 2.92) were significantly upregulated in Post-CP MIXO compared with Post-CP AUTO (Fig. 5E). Z-score profiling further showed enrichment of aspartate aminotransferase (*AST2*), proline dehydrogenase (*PDY1-2*), and threonine dehydratase (*THD1*). In line with this, several genes linked to branched-chain amino acid degradation were also elevated in Post-CP MIXO, including 3-hydroxyisobutyrate dehydrogenase (*HID1*), aldehyde dehydrogenase (*ALDH6*), and thiosulfate sulfurtransferase (*TSU1*), relative to Post-CP AUTO. Taken together, the depletion of branched-chain amino acids and proline supports enhanced amino acid catabolism and a concomitant rerouting of central carbon fluxes under MIXO. Proline catabolism is consistent with previous studies reporting upregulation of *PDY1–2*, indicating that proline oxidation can contribute to ATP production during nutrient remobilization in *Arabidopsis thaliana* (Launay et al., 2019). Likewise, acetylornithine deacetylase (*AOD1*; logFC = 2.85), ornithine aminotransferase (*OAT1*; logFC = 2.56), glutamate 5-kinase (*PROBI*; logFC = 2.49), and glycerol-3-phosphate dehydrogenase (GPD2; logFC = 2.83) were upregulated in Post-CP MIXO compared with Post-CP HETERO (Supplementary Dataset 5).

#### Effect of trophic regime on cell cycle regulation

In both AUTO and MIXO, *CDKA1* expression was relatively low in Pre-CP and increased in Post-CP (Fig. 7). The Z-score showed stronger *CDKA*1 upregulation in Post-CP MIXO than in Post-CP AUTO. *CDKB1* increased from Pre-CP to Post-CP in AUTO, but in MIXO it showed the opposite pattern, being reduced in Post-CP and relatively higher at Pre-CP. *CDKB1* expression has been associated first with pre-commitment and, second, much more strongly during the S/M phase (Bisova et al., 2005; Cross and Umen, 2015; Strenkert et al., 2019; Zones et al., 2015). Similar mitosis-associated upregulation has also been described for other CDKB-family members in plants (Magyar et al., 1997; Menges et al., 2002). *CDKC1* showed a similar pattern in both AUTO and MIXO, with lower expression at Pre-CP and higher expression at Post-CP (Fig. 6). *CDKC1* levels were consistently higher in MIXO than in AUTO at both Pre-CP and Post-CP. In contrast, *CDKD1* showed higher expression at Pre-CP than at Post-CP in both AUTO and MIXO, with overall expression again being higher under MIXO. *CYCA1* and *CYCB1* displayed patterns similar to *CDKB1*. The expression of *E2F1*–*DP1* and the retinoblastoma protein (*MAT3*) was higher at Post-CP than at Pre-CP under both AUTO and MIXO conditions; at the Post-CP stage, their expression was higher in MIXO than in AUTO (Olson et al., 2010).

**Figure 7.**
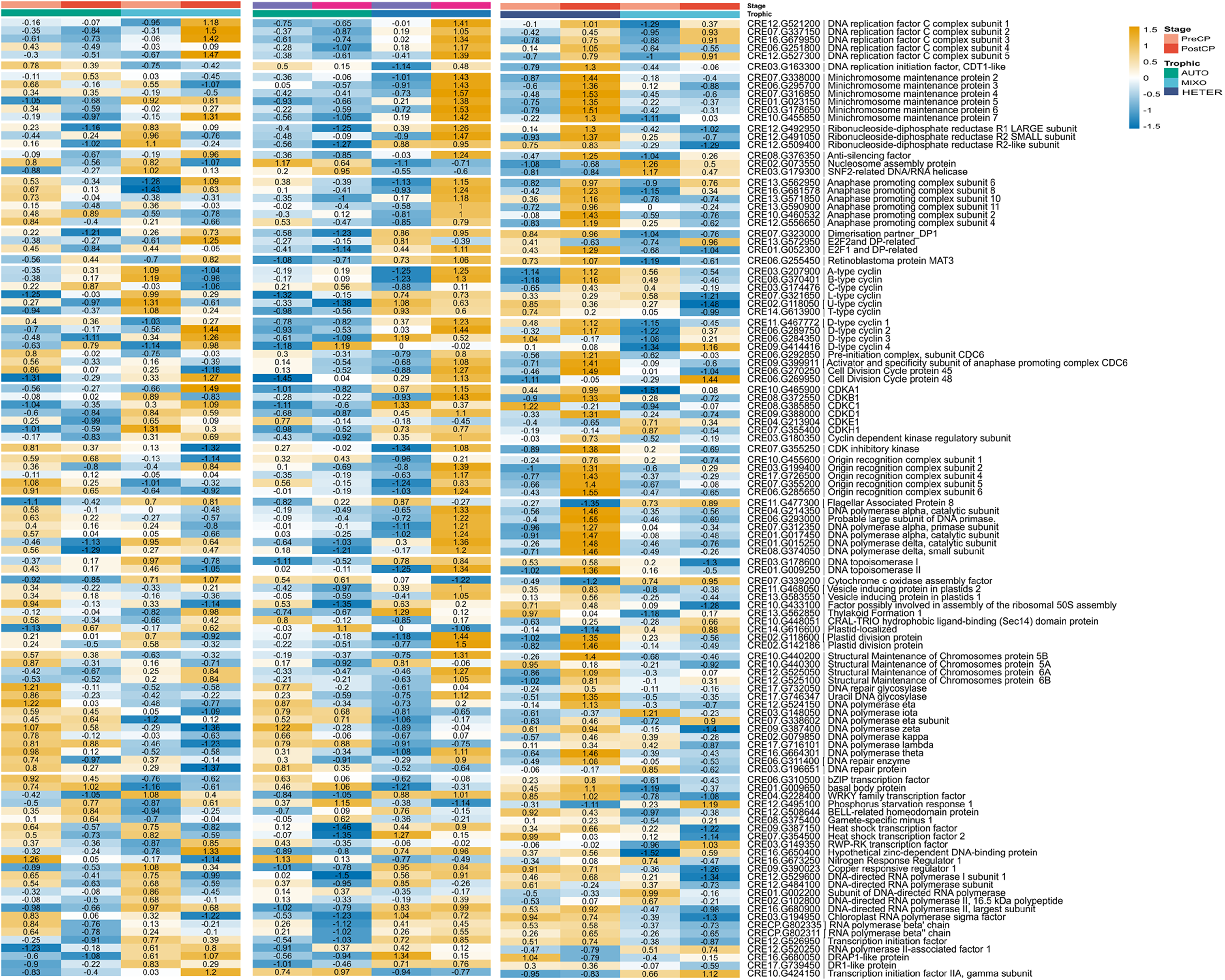
Z-score heatmaps show the expression of genes involved in the cell cycle in AUTO, MIXO, and HETERO under PreCP and Post-CP conditions. Colors indicate relative expression levels (yellow – increased, blue - decreased)

DNA polymerases *POLA2/4*, *PODA1/2*, and the ribonucleoside-diphosphate reductase large and small subunits (*RIR1–2*) showed higher expression at Pre-CP compared to Post-CP in both AUTO and MIXO. The *POLA2/4* and *PODA1/2* Pre-CP expression was higher in MIXO than in AUTO. Replication factor C subunits (*RFC1–5*) were strongly upregulated in Post-CP MIXO, together with increased expression of *MCM5–7*, whereas these DNA replication genes were downregulated in Post-CP AUTO, indicating active replication machinery in Post-CP MIXO (Supplementary Dataset 5).

When comparing MIXO and HETERO, *CDKA1* showed an increased expression trend from Pre-CP to Post-CP under both conditions; both PreCP and Post-CP expression levels were markedly higher in HETERO than in MIXO (Fig. 6). This is consistent with reports showing that CDKA1 protein is present throughout the cell cycle, whereas substantial CDKA1-associated kinase activity is detected primarily in actively dividing *C. reinhardtii* (Atkins and Cross, 2018). *CDKB1* showed higher expression in Pre-CP MIXO and was downregulated at Post-CP MIXO, whereas the opposite trend was observed in HETERO, with lower expression at Pre-CP and increased expression at Post-CP. CDKB1 is associated first with pre-commitment and is most strongly induced during S/M (Bisova et al., 2005; Cross and Umen, 2015; Strenkert et al., 2019; Zones et al., 2015). *CYCA1* and *CYCB1* showed higher expression at Post-CP in HETERO compared to MIXO.

Transcripts corresponding to the E2F1–DP1 regulatory module and the MAT3/RB complex were more abundant in Post-CP HETERO than in Post-CP MIXO. Consistently, Post-CP HETERO also exhibited markedly higher expression of core cell cycle and DNA replication machinery, including *APC1–10*, *MCM2–7*, and *RFC1–5*, relative to Post-CP MIXO. In addition, Post-CP HETERO showed elevated expression of genes implicated in DNA damage tolerance and repair, including DNA polymerase η (*POLH2*), DNA polymerase ι (*POLI1*), DNA polymerase ζ (*POLZ1*), and the structure-specific endonuclease *XPF1* (Fig. 6, Supplementary Dataset 5). Likewise, higher expression of *WEE1* was found in Post-CP HETERO than in Post-CP MIXO.

Wee1 kinase is regarded as a mitotic inhibitor and acts as a checkpoint for DNA damage response. In *Arabidopsis thaliana*, elevated *WEE1* kinase expression suppressed growth by arresting cells in the G phase of the cell cycle, acting as part of the DNA integrity checkpoint that couples mitosis to DNA repair in cells that suffer DNA damage (De Schutter et al., 2007).

## Discussion

### Growth and photosynthesis

Acetate supplementation is known to enhance metabolism and accelerate growth in many green algae (Lauersen et al., 2016; Rai et al., 2013; Smith et al., 2015). Similarly, there was greater accumulation of total lipid and starch in MIXO compared to AUTO and HETERO (Fig. 2h, k). This aligns with metabolite studies showing higher accumulation of both starch and triacylglycerol (TAG) in mixotrophy compared to heterotrophy (Singh et al., 2014). This mechanism appears to be general for other algae, such as *Parachlorella kessleri* TY (Gao et al., 2023). Light harvesting efficiency is influenced by both the composition and concentration of photosynthetic pigments. Pigment concentration depends on biomass density and growth conditions, while composition is primarily affected by the intensity and quality of light (Bialevich et al., 2022). Among all treatments, AUTO showed the highest total chlorophyll content, accompanied by high Fv/Fm. A higher Fv/Fm ratio is commonly regarded as increased photosynthetic efficacy, whereas a lower ratio implies PSII photoinhibition (Murata et al., 2007). The lowest chlorophyll content was found in HETERO; however, Fv/Fm remained stable. This is consistent with other algae, where total chlorophyll moderately decreases under mixotrophic conditions and is severely reduced in heterotrophy (Cheirsilp and Torpee, 2012).

Based on the transcriptomics, photosynthetic traits were not significantly affected in MIXO compared to AUTO. However, antenna subunits (LHCA4, LHCBM1, LHCII) and cytochrome b6/f complex genes (petD and petG) were upregulated in Post-CP MIXO. In contrast, the RuBisCO large subunit (rbcL) decreased slightly in MIXO, consistent with the established effect of acetate minimizing biosynthetic and energetic investment when an efficient external carbon source is available (Heifetz et al., 2000; Koley et al., 2026; Kroymann et al., 1995). Together, these data suggest that MIXO cells maintain photochemical electron transport (Wilken et al., 2013) while selectively reducing investment in carbon fixation machinery, consistent with acetate partially substituting for carbon derived from the CBB cycle (see below). Acetate-derived carbon can substitute for half the photoautotrophically generated biomass, while photosynthesis provides ATP and reductant for acetate assimilation (Eppley and MaciasR, 1963). In heterotrophy, the expression of the PSI light-harvesting component (LHCA4), the PSII core reaction-center genes (psbA (D1) and psbD (D2)), and the RuBisCO subunits (rbcL, RBCS1, and RBCS2) increased, consistent with observations in other green algae (Recuenco-Munoz et al., 2015; Zhang et al., 2021). This suggests that PSI/PSII and carbon-fixation machinery are maintained under heterotrophy, rather than being fully unused (Fig. 6). Whether this reflects functional maintenance or transcriptional persistence without corresponding activity is yet to be determined, but Fv/Fm ratios suggest that PSII is potentially functional.

### Acetate assimilation

Mixotrophy reorganizes central carbon metabolism to optimize carbon retention, redox balance, and biosynthetic fluxes. Acetate assimilation, or acetyl-CoA ligation, occurs to varying extents in all three intracellular compartments: 1) via a cytosolic citrate synthase followed by transport into mitochondria for the TCA cycle, 2) by direct uptake into mitochondria, conversion through a mitochondrial citrate synthase, and processing through the TCA cycle, and 3) by direct assimilation into fatty acids in the plastid (Boyle et al., 2017). Furthermore, acetate assimilation is associated with an increased number of peroxisomes and elevated expression of genes encoding putative glyoxysomal proteins (Hayashi et al., 2015). In our experiments under MIXO, both transcripts linked to acetate assimilation (GFY1–5, ACS1, PAT) and to glyoxylate shunt genes (ICL1, MSA1) were upregulated. However, these genes were expressed even more strongly in Post-CP HETERO, indicating that acetate assimilation and glyoxylate cycle flux were more active in HETERO than in MIXO. This is consistent with the central role of the glyoxylate cycle in acetate assimilation in both HETERO and MIXO (Plancke et al., 2014). In addition to the glyoxylate cycle, the TCA cycle remains active under mixotrophy, with acetate-derived carbon rerouted through the glyoxylate shunt to produce succinate and malate while bypassing TCA decarboxylation steps and conserving carbon (Koley et al., 2026). Consistent with this, nearly all TCA cycle transcripts were upregulated in Post-CP MIXO compared to Post-CP AUTO and Post-CP HETERO (Fig. 8).

**Figure 8.**
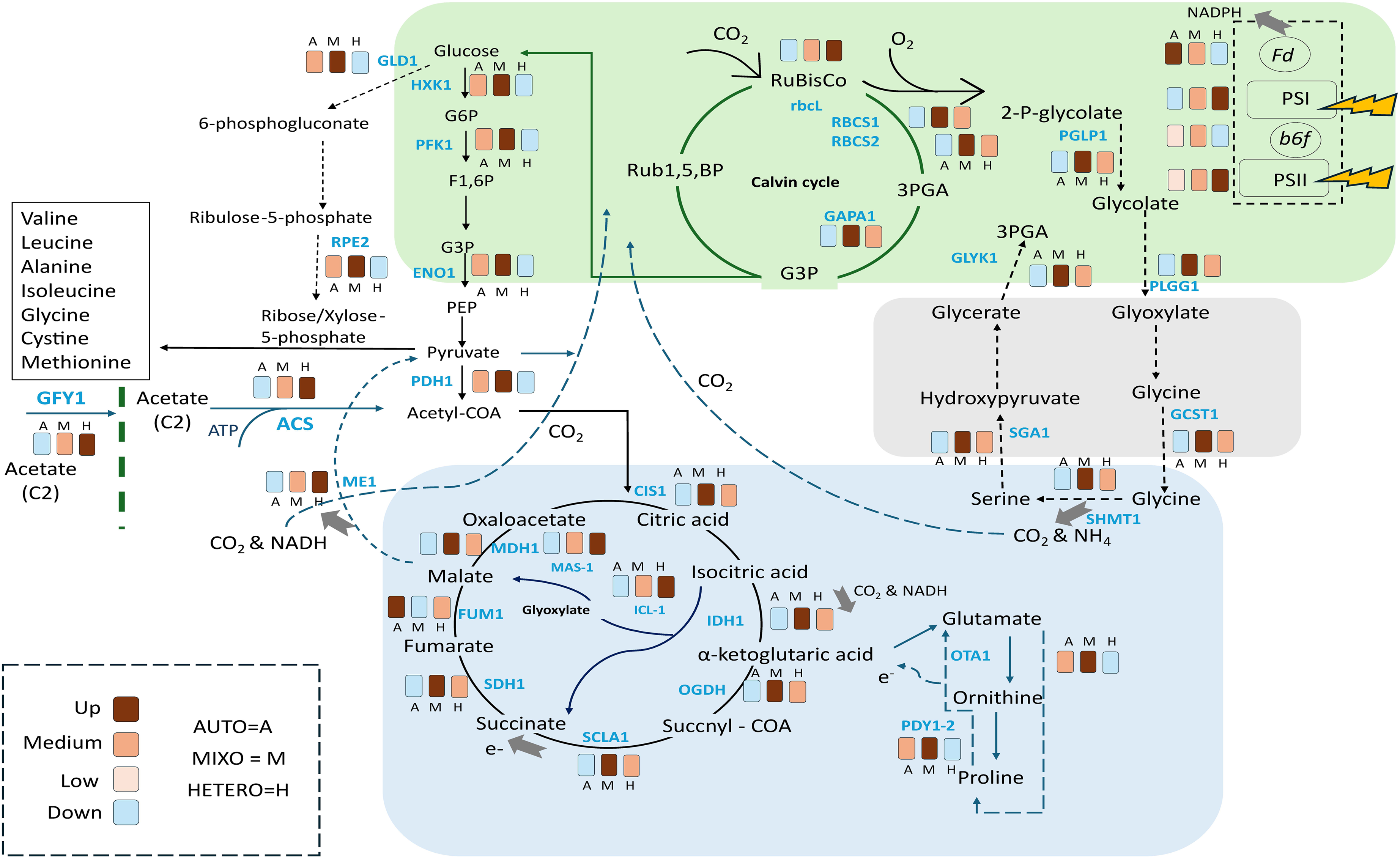
Systematic representation of metabolic changes under phototrophic, mixotrophic, and heterotrophic growth conditions in *Chlamydomonas reinhardtii*, with gene expression at Post-CP and CP time points shown here. *rbcL*: RuBisCO large subunit; *RBCS1*: ribulose-1,5-bisphosphate carboxylase/oxygenase small subunit 1, chloroplastic; *RBCS2*: ribulose-1,5-bisphosphate carboxylase/oxygenase small subunit 2, chloroplastic; *GAPA1*: glyceraldehyde-3-phosphate dehydrogenase A; *PGLP1*: phosphoglycolate phosphatase; *PLGG1*: plastidic glycolate/glycerate transporter; *SHMT1*: serine hydroxymethyltransferase; *SGA1*: glyoxylate aminotransferase; *GLYK*: glycerate kinase; *GLD1*: glucose-6-phosphate-1-dehydrogenase; *HXK1*: hexokinase; *PFK1*: phosphofructokinase; *ENO1*: Enolase; *PDH1*: Pyruvate dehydrogenase; *RPE2*: ribulose phosphate-3-epimerase; *CIS1*: citrate synthase, mitochondrial; *IDH1*: isocitrate dehydrogenase; *OGDH*: 2-oxoglutarate dehydrogenase; *SCLA1*: succinyl-CoA ligase; *SDH1*: succinate dehydrogenase; *FUM1*: fumarate hydratase; *MDH1*: malate dehydrogenase; *ME1*: malic enzyme; *ACS*: acetyl-CoA synthetase; *GFY*: putative acetate transporter; *OTA1*: ornithine transaminase; *PDY1*: proline dehydrogenase.

During mixotrophy in the green alga *Chlorella sorokiniana*, light enhances photorespiration to accelerate the glyoxylate cycle and release acetate-derived CO for the CBB cycle (Xie et al., 2015). Specifically, photorespiration produces glyoxylate, which, in the glyoxylate cycle, condenses with acetyl-CoA to form malate, effectively linking photorespiration with acetate assimilation. Glyoxylate can be converted into glycine, releasing CO from acetate, followed by carbon fixation in the CBB cycle. A similar mechanism may also occur in C. reinhardtii, as transcripts associated with photorespiration – glycolate transporter (PLGG1), glycine decarboxylase complex (GCST1), and serine hydroxymethyltransferase (SHMT1)—were upregulated in Post-CP MIXO (Fig. 8). The upregulation of GCST1 and SHMT1 indicates that glycine is transported into the mitochondria, where two glycine molecules undergo deamination and decarboxylation by GCST1 and SHMT1, resulting in the formation of one molecule each of serine, ammonia, and carbon dioxide (Pick et al., 2013). In HETERO, these transcripts were also upregulated, with abundance exceeding AUTO but remaining below MIXO, consistent with photorespiration being maintained in heterotrophy but operating at a reduced level compared with mixotrophy. In line with this, several CBB transcripts (RuBisCO subunits rbcL, RBCS1, RBCS2, fructose-1,6-bisphosphate aldolase (FBA), NADPH-dependent malate dehydrogenase (NADP-MDH)) were elevated in HETERO, suggesting continued CO fixation, with a contribution from internally released CO from acetate catabolism. Flux balance analysis suggested that mixotrophy also activates the oxidative pentose phosphate (OPP) pathway (Chapman et al., 2016), which functions as a carbon sink, redirecting photosynthate toward nucleotide biosynthesis and thereby supporting increased growth rate and biomass production under mixotrophy. In this way, mixotrophy supports mitochondrial respiration while reducing carbon-fixation flux and NADPH demand (Chapman et al., 2016). In our experiments, OPP pathway intermediates were most strongly upregulated in Post-CP MIXO, showed a more modest increase in Post-CP AUTO, and were downregulated in Post-CP HETERO (Fig. 8). The downregulation in HETERO may be consistent with an inactive OPP pathway in heterotrophy in another green alga, *Chlorella vulgaris* (Zuniga et al., 2016). Taken together, the transcriptional profile supports coordinated metabolic rewiring under mixotrophy, in which acetate assimilation, glyoxylate shunt activation, increased TCA flux, and elevated photorespiration collectively promote carbon conservation and redox flexibility. Rather than representing parallel pathways, these modules appear functionally integrated to maximize biomass production while minimizing carbon loss.

### Stress response

The strong transcriptional induction of stress-associated genes in HETERO likely reflects the metabolic imbalance resulting from reliance on acetate without photosynthetic energy input. The HETERO conditions rapidly induce transcripts that encode heat shock proteins (HSPs, HSP70), the ribosome-associated molecular co-chaperone (ZRF1), and reactive oxygen species (ROS) scavenging genes, including superoxide dismutases (FSD1, MSD1), glutathione peroxidases (GPX1/2/++), catalases (CAT1/2/3), and gamma-tocopherol methyltransferase (VTE4). HSP70 genes are naturally induced by a dark-to-light shift in C. reinhardtii (Schroda et al., 1999; von Gromoff et al., 1989) and are regulated by a circadian rhythm to prepare cells for subsequent exposure to high light (Kropat et al., 1995; Schuster et al., 1988). The HSPs (HSP90, HSP110, mitochondrial HSP70), genes involved in superoxide dismutase activity, glutathione- and thiol-based redox regulation, and tocopherol biosynthesis were also upregulated in MIXO, suggesting that, rather than indicating stress, this expression is related to metabolic reprogramming. This upregulation of HSP70 seems to be a general mechanism as it was shown also in mixotrophically grown *Chlorella vulgaris* (Arora and Philippidis, 2021). Thus, HSP70 induction may reflect primarily remodeling demands, such as rebalancing light-harvesting complexes, PSII repair, thylakoid protein turnover, and changes in ROS production during trophic transition or possible proteostatic stress.

### Cell cycle in different trophic regimes

The original hypothesis driving this study was to identify common and distinct transcriptional patterns between the trophic regimes. While the common features will be described elsewhere, here we focus on differences among the treatments. Comparing Post-CP MIXO with Post-CP AUTO showed similar patterns of cell cycle gene expression with generally higher expression in MIXO conditions (Fig. 7). We were able to confirm the established patterns (Bisova et al., 2005) of CDKB1, CYCA1, and CYCB1 with increase in Pre-CP MIXO and decreasing in Post-CP MIXO. Because our sampling was around the CP, we were unable to observe the major upregulation of these genes during the S/M phase (Bisova et al., 2005; Strenkert et al., 2019; Zones et al., 2015). In contrast, the Post-CP HETERO showed increased expression of most cell cycle genes, including those involved in DNA replication, DNA replication initiation factors, the MCM complex, CDKs, ribonucleosides, and plastid division (Fig. 7). The CDK-inhibitory kinase WEE1 was highly expressed in Post-CP HETERO, along with several DNA repair factors, including DNA repair glycosylase (AGE3), uracil DNA glycosylase (UNG1), and translesion DNA polymerases η, ι, ζ, and κ. Similarly, leucine zipper (bZIP) and WRKY family transcription factors were also upregulated in Post-CP HETERO. Both families are associated with stress-responsive regulation, with bZIPs reported in oxidative stress tolerance in C. reinhardtii (Choi et al., 2022; Ji et al., 2018) and WRKYs widely reported in plant stress responses (Arora and Philippidis, 2021; Boonyaves et al., 2022; Wang et al., 2024). A simple explanation stems from very slow growth and late CP attainment in HETERO, which is reflected in the sampling. For both AUTO and MIXO, the Post-CP samples were collected early in the cell cycle and at least six hours before the onset of S/M phase. In contrast, in HETERO, the Post-CP sample preceded the onset of S/M phase by only 2–3 hours. Therefore, it is plausible that the established upregulation of cell cycle genes, known to occur before S/M onset, had already begun in the Post-CP HETERO samples. Given that WEE1 transcription increases prior to S/M entry, its elevated expression in HETERO is consistent with a later sampling window closer to mitotic onset (Zones et al., 2015) rather than a stress-induced checkpoint activation. Alternatively, it remains possible that genes encoding proteins involved in cell division, especially CDKs, DNA replication, and mitotic entry, were abnormally and overwhelmingly more highly expressed in Post-CP CP HETERO compared to Post-CP CP MIXO and Post-CP CP AUTO. This correlates with the commitment point analysis (Fig. ---), which showed that fewer cells reached the commitment point in HETERO and that cell-cycle progression was delayed compared with the other conditions. This would be consistent with observations in the C. reinhardtii cht7 mutant grown under nitrogen-depleted conditions, which also shows higher expression of cell-cycle genes with ≥2C DNA and reduced growth, indicating premature or unscheduled DNA replication compared to wild type (Lin et al., 2022; Takeuchi et al., 2020).

Collectively, these findings suggest that trophic regime affects not only metabolic flux distribution but also redox balance, proteostasis, and the timing of cell-cycle transitions. Our results indicate that autotrophic, mixotrophic, and heterotrophic growth in Chlamydomonas are associated with distinct transcriptional adjustments that coordinate central carbon metabolism, photosynthetic capacity, and cell-cycle progression. Mixotrophy does not appear to represent a simple additive coexistence of autotrophic and heterotrophic programs; instead, it involves coordinated metabolic adjustments linked to altered carbon fixation investment and cell-cycle progression. In contrast, heterotrophy weakens the connection between metabolic input and cell-cycle timing, resulting in delayed commitment and stress-associated transcriptional signatures. These findings provide a framework for understanding how carbon availability influences metabolic organization and cell-cycle regulation in photosynthetic eukaryotes.

## Acknowledgements

We are obliged to the technical staff of the Laboratory of Cell Cycles of Algae for excellent technical support.

---

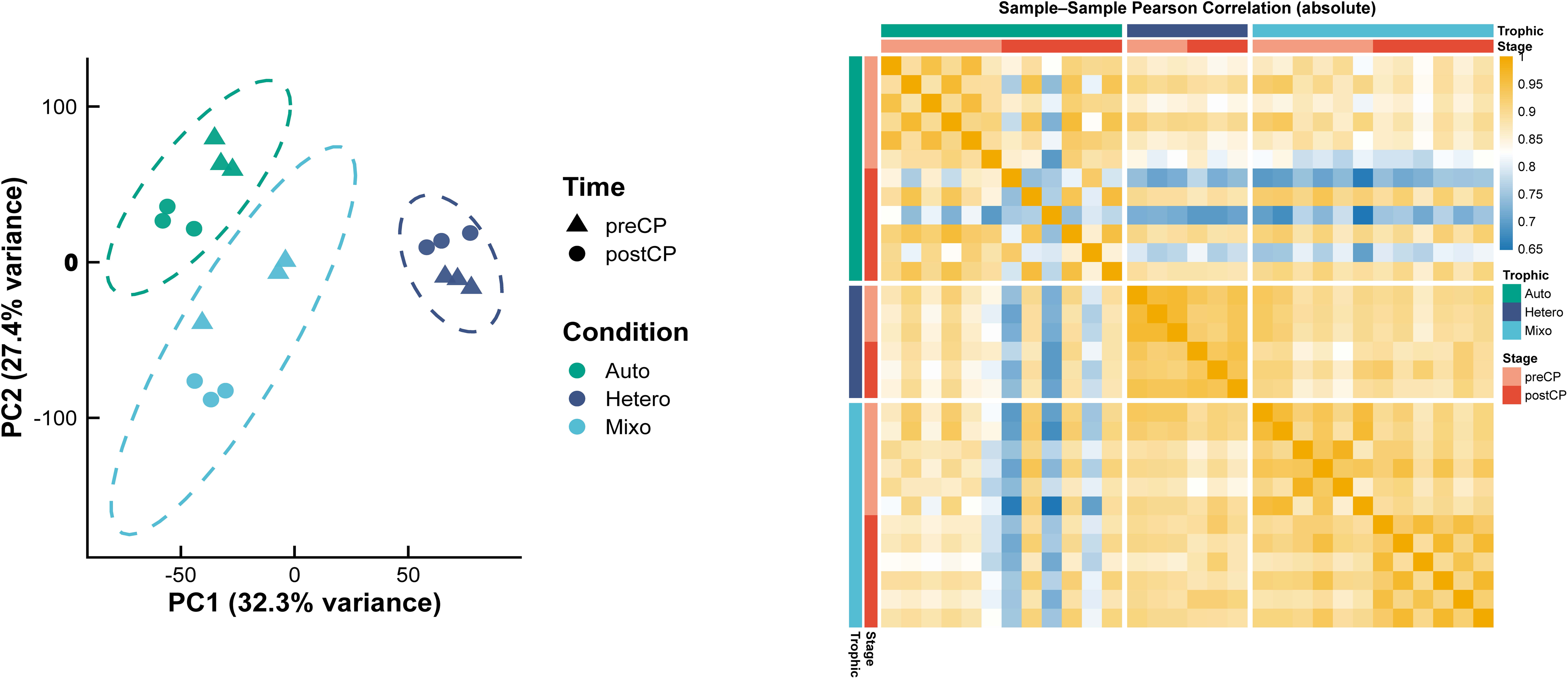

## References

Arora, N., and G.P. Philippidis. 2021. Unraveling metabolic alterations in *Chlorella vulgaris* cultivated on renewable sugars using time resolved multi-omics. Sci Total Environ 800:149504.

Atkins, K.C., and F. Cross. 2018. Inter-regulation of CDKA/CDK1 and the plant-specific cyclin-dependent kinase CDKB in control of the *Chlamydomonas* cell cycle. Plant Cell 30:429–446.

Baker, N.R. 2008. Chlorophyll fluorescence: a probe of photosynthesis in vivo. Annu Rev Plant Biol 59:89–113.

Bialevich, V., V. Zachleder, and K. Bišová. 2022. The effect of variable light source and light intensity on the growth of three algal species. Cells 11:1293.

Bisova, K., D.M. Krylov, and J.G. Umen. 2005. Genome-wide annotation and expression profiling of cell cycle regulatory genes in *Chlamydomonas reinhardtii*. Plant Physiol 137:475–491.

Bišová, K., and V. Zachleder. 2014. Cell-cycle regulation in green algae dividing by multiple fission. J Exp Bot 65:2585–2602.

Boonyaves, K., T.Y. Wu, Y. Dong, and D. Urano. 2022. Interplay between ARABIDOPSIS Gbeta and WRKY transcription factors differentiates environmental stress responses. Plant Physiol 190:813–827.

Boyle, N.R., and J.A. Morgan. 2009. Flux balance analysis of primary metabolism in Chlamydomonas reinhardtii. BMC Syst Biol 3:4.

Boyle, N.R., N. Sengupta, and J.A. Morgan. 2017. Metabolic flux analysis of heterotrophic growth in *Chlamydomonas reinhardtii*. PLoS One 12:e0177292.

Brányiková, I., B. Maršálková, J. Doucha, T. Brányik, K. Bišová, V. Zachleder, and M. Vítová. 2011. Microalgae-novel highly efficient starch producers. Biotechnol Bioeng 108:766–776.

Burgess, S.J., H. Taha, J.A. Yeoman, O. Iamshanova, K.X. Chan, M. Boehm, V. Behrends, J.G. Bundy, W. Bialek, J.W. Murray, and P.J. Nixon. 2016. Identification of the elusive pyruvate reductase of *Chlamydomonas reinhardtii* chloroplasts. Plant Cell Physiol 57:82–94.

Castaño-Cerezo, S., V. Bernal, H. Post, T. Fuhrer, S. Cappadona, N.C. Sánchez-Díaz, U. Sauer, A.J. Heck, A.F. Altelaar, and M. Cánovas. 2014. Protein acetylation affects acetate metabolism, motility and acid stress response in Escherichia coli. Mol Syst Biol 10:762.

Chapman, S.P., C.M. Paget, G.N. Johnson, and J.M. Schwartz. 2016. Corrigendum: Flux balance analysis reveals acetate metabolism modulates cyclic electron flow and alternative glycolytic pathways in *Chlamydomonas reinhardtii*. Front Plant Sci 7:362.

Cheirsilp, B., and S. Torpee. 2012. Enhanced growth and lipid production of microalgae under mixotrophic culture condition: effect of light intensity, glucose concentration and fed-batch cultivation. Bioresour Technol 110:510–516.

Chen, J., Y. Chen, W. He, H. Liang, T. Hong, T. Li, and H. Du. 2024. Transcriptome analysis reveals the molecular mechanism of differences in growth between photoautotrophy and heterotrophy in Chlamydomonas reinhardtii. Front Plant Sci Volume 15–2024:

Chen, S. 2023. Ultrafast one-pass FASTQ data preprocessing, quality control, and deduplication using fastp. iMeta 2:e107.

Chen, Y., L. Chen, A.T.L. Lun, P.L. Baldoni, and G.K. Smyth. 2025. edgeR v4: powerful differential analysis of sequencing data with expanded functionality and improved support for small counts and larger datasets. Nucleic Acids Res 53:

Choi, B.Y., H. Kim, D. Shim, S. Jang, Y. Yamaoka, S. Shin, T. Yamano, M. Kajikawa, E. Jin, H. Fukuzawa, and Y. Lee. 2022. The *Chlamydomonas* bZIP transcription factor BLZ8 confers oxidative stress tolerance by inducing the carbon-concentrating mechanism. Plant Cell 34:910–926.

Coleman, A.W. 1982. The nuclear-cell cycle in *Chlamydomonas* (Chlorophyceae). J Phycol 18:192–195.

Craig, R.J., S.D. Gallaher, S. Shu, P.A. Salomé, J.W. Jenkins, C.E. Blaby-Haas, S.O. Purvine, S. O’Donnell, K. Barry, J. Grimwood, D. Strenkert, J. Kropat, C. Daum, Y. Yoshinaga, D.M. Goodstein, O. Vallon, J. Schmutz, and S.S. Merchant. 2022. The Chlamydomonas Genome Project, version 6: Reference assemblies for mating-type plus and minus strains reveal extensive structural mutation in the laboratory. Plant Cell 35:644–672.

Cross, F.R., and J.G. Umen. 2015. The *Chlamydomonas* cell cycle. Plant J 82:370–392.

De Schutter, K., J. Joubes, T. Cools, A. Verkest, F. Corellou, E. Babiychuk, E. Van Der Schueren, T. Beeckman, S. Kushnir, D. Inze, and L. De Veylder. 2007. Arabidopsis WEE1 kinase controls cell cycle arrest in response to activation of the DNA integrity checkpoint. Plant Cell 19:211–225.

Decallonne, J.R., and J.C. Weyns. 1976. A shortened procedure of the diphenylamine reaction for the measurement of deoxyribonucleic acid by using light activation. Anal Biochem 74:448–456.

Dupuis, S., and S.S. Merchant. 2023. *Chlamydomonas reinhardtii*: a model for photosynthesis and so much more. Nat Methods 20:1441–1442.

Durante, L., W. Hubner, K.J. Lauersen, and C. Remacle. 2019. Characterization of the GPR1/FUN34/YaaH protein family in the green microalga *Chlamydomonas* suggests their role as intracellular membrane acetate channels. Plant Direct 3:e00148.

Endo, T., and K. Asada. 1996. Dark induction of the non-photochemical quenching of chlorophyll fluorescence by acetate in *Chlamydomonas reinhardtii*. Plant Cell Physiol 37:551–555.

Eppley, R.W., and F.M. MaciasR. 1963. Role of the alga *Chlamydomonas mundana* in anaerobic waste stabilization lagoons. Limnol. Oceanogr. 8:411–416.

Fauser, F., J. Vilarrasa-Blasi, M. Onishi, S. Ramundo, W. Patena, M. Millican, J. Osaki, C. Philp, M. Nemeth, P.A. Salome, X. Li, S. Wakao, R.G. Kim, Y. Kaye, A.R. Grossman, K.K. Niyogi, S.S. Merchant, S.R. Cutler, P. Walter, J.R. Dinneny, M.C. Jonikas, and R.E. Jinkerson. 2022. Systematic characterization of gene function in the photosynthetic alga *Chlamydomonas reinhardtii*. Nat Genet 54:705–714.

Fett, J.P., and J.R. Coleman. 1994. Regulation of periplasmic carbonic anhydrase expression in *Chlamydomonas reinhardtii* by acetate and pH. Plant Physiol 106:103–108.

Füßl, M., A.-C. König, J. Eirich, M. Hartl, L. Kleinknecht, A.-V. Bohne, A. Harzen, K. Kramer, D. Leister, J. Nickelsen, and I. Finkemeier. 2022. Dynamic light- and acetate-dependent regulation of the proteome and lysine acetylome of *Chlamydomonas*. Plant J 109:261–277.

Gao, Y., Y. Li, Y. Yang, J. Feng, L. Ji, and S. Xie. 2023. Effects of trophic modes on the lipid accumulation of *Parachlorella kessleri* TY. Fermentation 9:891.

Genty, B., J.-M. Briantais, and N.R. Baker. 1989. The relationship between the quantum yield of photosynthetic electron transport and quenching of chlorophyll fluorescence. Biochim Biophys Acta - General Subjects 990:87–92.

Gorman, D.S., and R.P. Levine. 1965. Cytochrome f and plastocyanin: their sequence in the photosynthetic electron transport chain of *Chlamydomonas reinhardi*. Proc Natl Acad Sci U S A 54:1665–1669.

Guerrero, S., V. Roces, L. García-Campa, L. Valledor, and M. Meijón. 2024. Proteomic dynamics revealed sex-biased responses to combined heat-drought stress in *Marchantia*. J Integr Plant Biol 66:2226–2241.

Harris, E.H. 2001. *Chlamydomonas* as a model organism. Annu Rev Plant Physiol Plant Mol Biol 52:363–406.

Hayashi, Y., N. Sato, A. Shinozaki, and M. Watanabe. 2015. Increase in peroxisome number and the gene expression of putative glyoxysomal enzymes in *Chlamydomonas* cells supplemented with acetate. Journal of Plant Research 128:177–185.

Heifetz, P.B., B. Förster, C.B. Osmond, L.J. Giles, and J.E. Boynton. 2000. Effects of acetate on facultative autotrophy in *Chlamydomonas reinhardtii* assessed by photosynthetic measurements and stable isotope analyses. Plant Physiol 122:1439–1446.

Hlavová, M., M. Vítová, and K. Bišová. 2016. Synchronization of green algae by light and dark regimes for cell cycle and cell division studies. In Plant Cell Division. M.-C. Caillaud, editor Springer Science, New York, Heilderberg, Dordrecht, London. 3–16.

Hoober, J.K. 1989. The Chlamydomonas Sourcebook. A Comprehensive Guide to Biology and Laboratory Use. Elizabeth H. Harris. Academic Press, San Diego, CA, 1989. xiv, 780 pp., illus. $145. Science 246:1503–1504.

Ji, C., X. Mao, J. Hao, X. Wang, J. Xue, H. Cui, and R. Li. 2018. Analysis of bZIP transcription factor family and their expressions under salt stress in *Chlamydomonas reinhardtii*. Int J Mol Sci 19:

Johnson, X., and J. Alric. 2013. Central carbon metabolism and electron transport in *Chlamydomonas reinhardtii*: Metabolic constraints for carbon partitioning between oil and starch. Eukaryot Cell 12:776–793.

Kaur, S., J. Hérault, A. Caruso, G. Pencréac’h, M. Come, L. Gauvry, S. Claverol, and C. Loiseau. 2021. Proteomics and expression studies on lipids and fatty acids metabolic genes in *Isochrysis galbana* under the combined influence of nitrogen starvation and sodium acetate supplementation. Bioresource Technology Reports 15:100714.

Kindle, K.L. 1987. Expression of a gene for a light-harvesting chlorophyll a/b-binding protein in *Chlamydomonas reinhardtii*: effect of light and acetate. Plant Mol Biol 9:547–563.

Koley, S., K. Foley, Z. Perrine, S.A. Morley, S. Kambhampati, O. Gomez, K.L. Chu, Y.H. Chou, M. Wei, S.C. Tzeng, R. Williams, J.G. Umen, and D.K. Allen. 2026. Metabolic rewiring and biomass redistribution enable optimized mixotrophic growth in *Chlamydomonas*. Proc Natl Acad Sci U S A 123:e2522572123.

Kropat, J., E.D. von Gromoff, F.W. Müller, and C.F. Beck. 1995. Heat shock and light activation of a *Chlamydomonas* HSP70 gene are mediated by independent regulatory pathways. Mol Gen Genet 248:727–734.

Kroymann, J., W. Schneider, and K. Zetsche. 1995. Opposite regulation of the copy number and the expression of plastid and mitochondrial genes by light and acetate in the green flagellate *Chlorogonium*. Plant Physiol 108:1641–1646.

Kruger, N.J., and A. von Schaewen. 2003. The oxidative pentose phosphate pathway: structure and organisation. Curr Opin Plant Biol 6:236–246.

Kselíková, V., V. Zachleder, and K. Bišová. 2022. Analysis of commitment point attainment in algae dividing by multiple fission. In Plant Cell Division. M.-C. Caillaud, editor Springer US, 89–101.

Lauersen, K.J., R. Willamme, N. Coosemans, M. Joris, O. Kruse, and C. Remacle. 2016. Peroxisomal microbodies are at the crossroads of acetate assimilation in the green microalga *Chlamydomonas reinhardtii*. Algal Res 16:266–274.

Launay, A., C. Cabassa-Hourton, H. Eubel, R. Maldiney, A. Guivarc’h, E. Crilat, S. Planchais, J. Lacoste, M. Bordenave-Jacquemin, G. Clement, L. Richard, P. Carol, H.P. Braun, S. Lebreton, and A. Savoure. 2019. Proline oxidation fuels mitochondrial respiration during dark-induced leaf senescence in *Arabidopsis thaliana*. J Exp Bot 70:6203–6214.

Lin, Y.T., T. Takeuchi, B. Youk, J. Umen, B.B. Sears, and C. Benning. 2022. Chlamydomonas CHT7 is involved in repressing DNA replication and mitotic genes during synchronous growth. G3 (Bethesda) 12:

Liu, D., C.A. Vargas-Garcia, A. Singh, and J. Umen. 2023. A cell-based model for size control in the multiple fission alga *Chlamydomonas reinhardtii*. Curr Biol 33:5215–5224 e5215.

Love, M.I., W. Huber, and S. Anders. 2014. Moderated estimation of fold change and dispersion for RNA-seq data with DESeq2. Genome Biol 15:550.

Lukavský, J., K. Tetík, and J. Vendlová. 1973. Extraction of nucleic acid from the alga *Scenedesmus quadricauda.* Archive für Hydrobiologie, Supplement 41, Algological Studies 9:416–426.

Magyar, Z., T. Meszaros, P. Miskolczi, M. Deak, A. Feher, S. Brown, E. Kondorosi, A. Athanasiadis, S. Pongor, M. Bilgin, L. Bako, C. Koncz, and D. Dudits. 1997. Cell cycle phase specificity of putative cyclin-dependent kinase variants in synchronized alfalfa cells. Plant Cell 9:223–235.

Martínez-Rivas, J.M., and J.M. Vega. 1993. Effect of culture conditions on the isocitrate dehydrogenase and isocitrate lyase activities in Chlamydomonas reinhardtii. Physiol Plant 88:599–603.

Menges, M., L. Hennig, W. Gruissem, and J.A. Murray. 2002. Cell cycle-regulated gene expression in Arabidopsis. J Biol Chem 277:41987–42002.

Merchant, S.S., S.E. Prochnik, O. Vallon, E.H. Harris, S.J. Karpowicz, G.B. Witman, A. Terry, A. Salamov, L.K. Fritz-Laylin, L. Marechal-Drouard, W.F. Marshall, L.-H. Qu, D.R. Nelson, A.A. Sanderfoot, M.H. Spalding, V.V. Kapitonov, Q. Ren, P. Ferris, E. Lindquist, H. Shapiro, S.M. Lucas, J. Grimwood, J. Schmutz, P. Cardol, H. Cerutti, G. Chanfreau, C.-L. Chen, V. Cognat, M.T. Croft, R. Dent, S. Dutcher, E. Fernandez, H. Fukuzawa, D. Gonzalez-Ballester, D. Gonzalez-Halphen, A. Hallmann, M. Hanikenne, M. Hippler, W. Inwood, K. Jabbari, M. Kalanon, R. Kuras, P.A. Lefebvre, S.D. Lemaire, A.V. Lobanov, M. Lohr, A. Manuell, I. Meier, L. Mets, M. Mittag, T. Mittelmeier, J.V. Moroney, J. Moseley, C. Napoli, A.M. Nedelcu, K. Niyogi, S.V. Novoselov, I.T. Paulsen, G. Pazour, S. Purton, J.-P. Ral, D.M. Riano-Pachon, W. Riekhof, L. Rymarquis, M. Schroda, D. Stern, J. Umen, R. Willows, N. Wilson, S.L. Zimmer, J. Allmer, J. Balk, K. Bisova, C.-J. Chen, M. Elias, K. Gendler, C. Hauser, M.R. Lamb, H. Ledford, J.C. Long, J. Minagawa, M.D. Page, J. Pan, W. Pootakham, S. Roje, A. Rose, E. Stahlberg, A.M. Terauchi, P. Yang, S. Ball, C. Bowler, C.L. Dieckmann, V.N. Gladyshev, P. Green, R. Jorgensen, S. Mayfield, B. Mueller-Roeber, S. Rajamani, R.T. Sayre, P. Brokstein, I. Dubchak, D. Goodstein, L. Hornick, Y.W. Huang, J. Jhaveri, Y. Luo, D. Martinez, W.C.A. Ngau, B. Otillar, A. Poliakov, A. Porter, L. Szajkowski, G. Werner, K. Zhou, I.V. Grigoriev, D.S. Rokhsar, and A.R. Grossman. 2007. The *Chlamydomonas* genome reveals the evolution of key animal and plant functions. Science 318:245–250.

Mielczarek, O., C.H. Rogers, Y. Zhan, L.S. Matheson, M.J.T. Stubbington, S. Schoenfelder, D.J. Bolland, B.M. Javierre, S.W. Wingett, C. Várnai, A. Segonds-Pichon, S.J. Conn, F. Krueger, S. Andrews, P. Fraser, L. Giorgetti, and A.E. Corcoran. 2023. Intra- and interchromosomal contact mapping reveals the Igh locus has extensive conformational heterogeneity and interacts with B-lineage genes. Cell Rep 42:113074.

Minhas, A.K., P. Hodgson, C.J. Barrow, and A. A. 2016. A review on the assessment of stress conditions for simultaneous production of microalgal lipids and carotenoids. Front Microbiol 7:1–19.

Murata, N., S. Takahashi, Y. Nishiyama, and S.I. Allakhverdiev. 2007. Photoinhibition of photosystem II under environmental stress. Biochim Biophys Acta - Bioenergetics 1767:414–421.

Neumann, G., R. Teras, L. Monson, M. Kivisaar, F. Schauer, and H.J. Heipieper. 2004. Simultaneous degradation of atrazine and phenol by *Pseudomonas sp.* strain ADP: effects of toxicity and adaptation. Appl Environ Microbiol 70:1907–1912.

Olson, B.J.S.C., M. Oberholzer, Y.B. Li, J.M. Zones, H.S. Kohli, K. Bisova, S.C. Fang, J. Meisenhelder, T. Hunter, and J.G. Umen. 2010. Regulation of the *Chlamydomonas* cell cycle by a stable, chromatin-associated retinoblastoma tumor suppressor complex. Plant Cell 22:3331–3347.

Paiva, S., F. Devaux, S. Barbosa, C. Jacq, and M. Casal. 2004. Ady2p is essential for the acetate permease activity in the yeast *Saccharomyces cerevisiae*. Yeast 21:201–210.

Patro, R., G. Duggal, M.I. Love, R.A. Irizarry, and C. Kingsford. 2017. Salmon provides fast and bias-aware quantification of transcript expression. Nat Methods 14:417–419.

Pick, T.R., A. Bräutigam, M.A. Schulz, T. Obata, A.R. Fernie, and A.P. Weber. 2013. PLGG1, a plastidic glycolate glycerate transporter, is required for photorespiration and defines a unique class of metabolite transporters. Proc Natl Acad Sci U S A 110:3185–3190.

Plancke, C., H. Vigeolas, R. Hohner, S. Roberty, B. Emonds-Alt, V. Larosa, R. Willamme, F. Duby, D. Onga Dhali, P. Thonart, S. Hiligsmann, F. Franck, G. Eppe, P. Cardol, M. Hippler, and C. Remacle. 2014. Lack of isocitrate lyase in *Chlamydomonas* leads to changes in carbon metabolism and in the response to oxidative stress under mixotrophic growth. Plant J 77:404–417.

Qiu, B., B. Xia, Q. Zhou, Y. Lu, M. He, K. Hasegawa, Z. Ma, F. Zhang, L. Gu, Q. Mao, F. Wang, S. Zhao, Z. Gao, and J. Liao. 2018. Succinate-acetate permease from *Citrobacter koseri* is an anion channel that unidirectionally translocates acetate. Cell Res 28:644–654.

Rai, M.P., S. Nigam, and R. Sharma. 2013. Response of growth and fatty acid compositions of *Chlorella pyrenoidosa* under mixotrophic cultivation with acetate and glycerol for bioenergy application. Biomass Bioenergy 58:251–257.

Recuenco-Munoz, L., P. Offre, L. Valledor, D. Lyon, W. Weckwerth, and S. Wienkoop. 2015. Targeted quantitative analysis of a diurnal RuBisCO subunit expression and translation profile in *Chlamydomonas reinhardtii* introducing a novel Mass Western approach. J Proteomics 113:143–153.

Roach, T., A. Sedoud, and A. Krieger-Liszkay. 2013. Acetate in mixotrophic growth medium affects photosystem II in *Chlamydomonas reinhardtii* and protects against photoinhibition. Biochim Biophys Acta 1827:1183–1190.

Robellet, X., M. Flipphi, S. Pégot, Andrew P. MacCabe, and C. Vélot. 2008. AcpA, a member of the GPR1/FUN34/YaaH membrane protein family, is essential for acetate permease activity in the hyphal fungus *Aspergillus nidulans*. Biochem J 412:485–493.

Sá-Pessoa, J., S. Paiva, D. Ribas, Inês J. Silva, Sandra C. Viegas, Cecília M. Arraiano, and M. Casal. 2013. SATP (YaaH), a succinate–acetate transporter protein in *Escherichia coli*. Biochem J 454:585–595.

Schroda, M., O. Vallon, F.A. Wollman, and C.F. Beck. 1999. A chloroplast-targeted heat shock protein 70 (HSP70) contributes to the photoprotection and repair of photosystem II during and after photoinhibition. Plant Cell 11:1165–1178.

Schuster, G., D. Even, K. Kloppstech, and I. Ohad. 1988. Evidence for protection by heat-shock proteins against photoinhibition during heat-shock. EMBO J 7:1–6.

Singh, H., M.R. Shukla, K.V. Chary, and B.J. Rao. 2014. Acetate and bicarbonate assimilation and metabolite formation in *Chlamydomonas reinhardtii*: a ^13^C-NMR study. PLoS One 9:e106457.

Smith, R.T., K. Bangert, S.J. Wilkinson, and D.J. Gilmour. 2015. Synergistic carbon metabolism in a fast growing mixotrophic freshwater microalgal species *Micractinium inermum*. Biomass Bioenergy 82:73–86.

Strenkert, D., S. Schmollinger, S.D. Gallaher, P.A. Salome, S.O. Purvine, C.D. Nicora, T. Mettler-Altmann, E. Soubeyrand, A.P.M. Weber, M.S. Lipton, G.J. Basset, and S.S. Merchant. 2019. Multiomics resolution of molecular events during a day in the life of *Chlamydomonas*. Proc Natl Acad Sci U S A

Takeuchi, T., B.B. Sears, C. Lindeboom, Y.T. Lin, N. Fekaris, K. Zienkiewicz, A. Zienkiewicz, E. Poliner, and C. Benning. 2020. *Chlamydomonas* CHT7 is required for an effective quiescent state by regulating nutrient-responsive cell cycle gene expression. Plant Cell 32:1240–1269.

Usadel, B., F. Poree, A. Nagel, M. Lohse, A. Czedik-Eysenberg, and M. Stitt. 2009. A guide to using MapMan to visualize and compare omics data in plants: a case study in the crop species, maize. Plant Cell Environ 32:1211–1229.

Vítová, M., K. Bišová, D. Umysová, M. Hlavová, S. Kawano, V. Zachleder, and M. Čížková. 2011. *Chlamydomonas reinhardtii*: duration of its cell cycle and phases at growth rates affected by light intensity. Planta 233:75–86.

von Gromoff, E.D., U. Treier, and C.F. Beck. 1989. Three light-inducible heat shock genes of *Chlamydomonas reinhardtii*. Mol Cell Biol 9:3911–3918.

Wang, W., H. Cao, J. Wang, and H. Zhang. 2024. Recent advances in functional assays of WRKY transcription factors in plant immunity against pathogens. Front Plant Sci 15:1517595.

Wanka, F. 1962. Die bestimmung der nucleinsäuren in *Chlorella pyrenoidosa*. Planta 58:594–619.

Weger, H.G., A.R. Chadderton, M. Lin, R.D. Guy, and D.H. Turpin. 1990. Cytochrome and alternative pathway respiration during transient ammonium assimilation by N-limited *Chlamydomonas reinhardtii*. Plant Physiol 94:1131–1136.

Wilken, S., J. Huisman, S. Naus-Wiezer, and E. Van Donk. 2013. Mixotrophic organisms become more heterotrophic with rising temperature. Ecol Lett 16:225–233.

Xie, X., A. Huang, W. Gu, Z. Zang, G. Pan, S. Gao, L. He, B. Zhang, J. Niu, A. Lin, and G. Wang. 2015. Photorespiration participates in the assimilation of acetate in *Chlorella sorokiniana* under high light. New Phytologist n/a–n/a.

Zachleder, V. 1984. Optimization of nucleic acids assay in green and blue-green algae: Extraction procedures and the light-activated reaction for DNA. Archive für Hydrobiologie, Supplement 67, Algological Studies 36:313–328.

Zhang, H., L. Zhao, Y. Chen, M. Zhu, Q. Xu, M. Wu, D. Han, and Q. Hu. 2021. Trophic transition enhanced biomass and lipid production of the unicellular green alga *Scenedesmus acuminatus*. Front Bioeng Biotechnol 9:638726.

Zhou, Y., F. Wu, J. Wu, S. Overmans, M. Ye, M. Xiao, B. Peng, L. Xu, J. Huang, Y. Lu, Y. Wang, S. Liang, H. Zhang, X. Liang, Z. Zhong, H. Liu, Z. Ruan, J. Xia, and P. Jin. 2024. The adaptive mechanisms of the marine diatom *Thalassiosira weissflogii* to long-term high CO_2_ and warming. Plant J 119:2001–2020.

Zones, J.M., I.K. Blaby, S.S. Merchant, and J.G. Umen. 2015. High-resolution profiling of a synchronized diurnal transcriptome from *Chlamydomonas reinhardtii* reveals continuous cell and metabolic differentiation. Plant Cell 10:2743–2769.

Zuniga, C., C.T. Li, T. Huelsman, J. Levering, D.C. Zielinski, B.O. McConnell, C.P. Long, E.P. Knoshaug, M.T. Guarnieri, M.R. Antoniewicz, M.J. Betenbaugh, and K. Zengler. 2016. Genome-scale metabolic model for the green alga *Chlorella vulgaris* UTEX 395 accurately predicts phenotypes under autotrophic, heterotrophic, and mixotrophic growth conditions. Plant Physiol

